# Conserved and Lineage-Specific Roles of KEA-Mediated Ion Homeostasis in *Chlamydomonas*

**DOI:** 10.64898/2025.12.03.692059

**Authors:** Tobias Wunder, Leon Eulitz, Luis Kramer, Zahin Mohd Ali, Matthias Ostermeier, Charlott Leu, Beata Szulc, Lorenz J. Holzner, Jonas Fechter, Francesco Padovani, Benjamin Brandt, Philipp Girr, Jing Tsong Teh, Susanne Mühlbauer, Clara Sotos, Max Angstenberger, Luke C. M. Mackinder, Kurt M. Schmoller, Joachim O. Rädler, Jörg Nickelsen, Jan de Vries, Hans-Henning Kunz

**Affiliations:** Plant Biochemistry and Physiology, LMU München, Großhaderner Straße 2-4, 82152, Martinsried-Planegg, Germany; Institute of Microbiology and Genetics, Göttingen Center for Molecular Biosciences (GZMB), Campus Institute Data Science (CIDAS), Department of Applied Bioinformatics, University of Göttingen, Goldschmidtstr. 1, 37077 Göttingen, Germany; Molecular Plant Science, LMU Munich, Großhaderner Straße 2-4, 82152 Planegg-Martinsried, Germany; Faculty of Physics and Center for NanoScience, Ludwig-Maximilian-University, Geschwister-Scholl-Platz 1, D-80539 Munich, Germany; Institute of Functional Epigenetics, Molecular Targets and Therapeutics Center, Helmholtz Zentrum Muenchen, 85764 Neuherberg, Germany; Centre for Novel Agricultural Products (CNAP), Department of Biology, University of York, Heslington, York YO10 5DD, UK; Institute for Integrative Biology of the Cell (I2BC), Université Paris-Saclay, CEA, CNRS, 91198 Gif-sur-Yvette cedex, France; Institute of Molecular Biosciences, Goethe University Frankfurt, 60438 Frankfurt, Germany

**Author notes:** **Correspondence:** Tobias Wunder Plant Biochemistry and Physiology LMU Munich Großhaderner Str. 2-4, 82152 Planegg-Martinsried, Germany. Senior authors: Tobias Wunder, Hans-Henning Kunz.

**Keywords:** photosynthesis, ion balance, rRNA processing, cell cycle, microalgae, chloroplast

## Abstract

In *Arabidopsis thaliana*, seamless plastid gene expression and development depend on finely balanced ion homeostasis across the inner envelope (IE) membrane, maintained by the K⁺/H⁺ antiporters AtKEA1/2. To assess whether these functions are retained across mono- and polyplastidic representatives of the green lineage, we studied CrKEA1, the sole IE KEA homolog in the unicellular alga *Chlamydomonas reinhardtii*. Using CRISPR/Cas9, we generated a *Cr-kea1* knockout mutant that exhibits impaired photoautotrophic growth, chloroplast deformation, and photoinhibition. Transcriptomics revealed strong induction of ribosome biogenesis genes and reduced abundance of transcripts associated with cell and plastid division. Further RNA analyses confirmed defects in stromal rRNA maturation of *Cr-kea1*, paralleling observations from Arabidopsis *kea1kea2* mutants. Expression of *CrKEA1* in Arabidopsis rescued growth and rRNA maturation in *At-kea1kea2*, demonstrating functional continuity after the ancient divergence between the two lineages. Cross-species transcriptomic comparisons further revealed that IE *KEA* loss elicits both shared and species-specific transcriptional responses: *PhANG* repression was conserved between algae and plants, whereas activation of the chloroplast unfolded protein response (cpUPR) and reduced expression of genes tied to cell-cycle and plastid fission occurred only in Chlamydomonas. Single-cell time-lapse imaging confirmed that *Cr-kea1* exhibits an increased frequency of aberrant cytokinesis, unequal division, and division failure. Our findings demonstrate that while IE KEA transporters fulfill conserved roles in maintaining the conditions for plastid gene expression, their integration into broader cellular networks has diverged between unicellular chlorophytes and embryophytes (land plants). This underscores a lineage-dependent tuning of plastid-nucleus communication shaped by organismal complexity and plastid number.

**One-sentence summary:** Disruption of KEA-mediated chloroplast ion homeostasis in *Chlamydomonas reinhardtii* reveals conserved and lineage-specific control of plastid rRNA processing and cell cycle progression.

## Introduction

A balanced ionic milieu is essential for plastid biogenesis and function (Kunz et al., 2024). In Arabidopsis (*Arabidopsis thaliana)*, disruption of the inner envelope (IE) K⁺/H⁺ antiporters AtKEA1 and AtKEA2 produces a virescent phenotype i.e., pale young leaves that eventually develop into green photosynthesizing tissue (Kunz et al., 2014). Subsequent work revealed that the essential role of AtKEA1/2 in plastid integrity and photosynthetic performance is linked to plastid gene expression and rRNA maturation (DeTar et al., 2021). These findings demonstrate a strong connection between organellar ion homeostasis and chloroplast biogenesis.

Dissecting this link mechanistically in plant cells is complicated due to their multicellular organization and polyplastidy: Plants typically harbor cells with dozens of plastids displaying heterogeneous behavior depending on tissue context and developmental stage (Knoblauch et al., 2024; Renna et al., 2025). This cellular heterogeneity makes it challenging to distinguish direct, plastid-autonomous consequences of ion imbalance from secondary effects at the tissue or organismal level.

Interestingly, many unicellular algae maintain a single plastid per cell. de Vries and Gould, 2018 described the evolutionary framework of the “monoplastidic bottleneck”, which posits that the ancestor of all Archaeplastida (Figure 1A) passed through a stage where plastid and host cell division were strictly synchronized. The unicellular green algae Chlamydomonas (*Chlamydomonas reinhardtii*), with its single cup-shaped chloroplast, represents a bona fide example of this ancestral state. In contrast, land plants such as Arabidopsis, have since evolved polyplastidy and decoupled plastid division from the nuclear cycle. Thus, Chlamydomonas is an ideal model to study the direct relation between plastid physiology, cell growth, and division, as each cell depends on a single plastid for both metabolism and successful cytokinesis.

**Figure 1:**
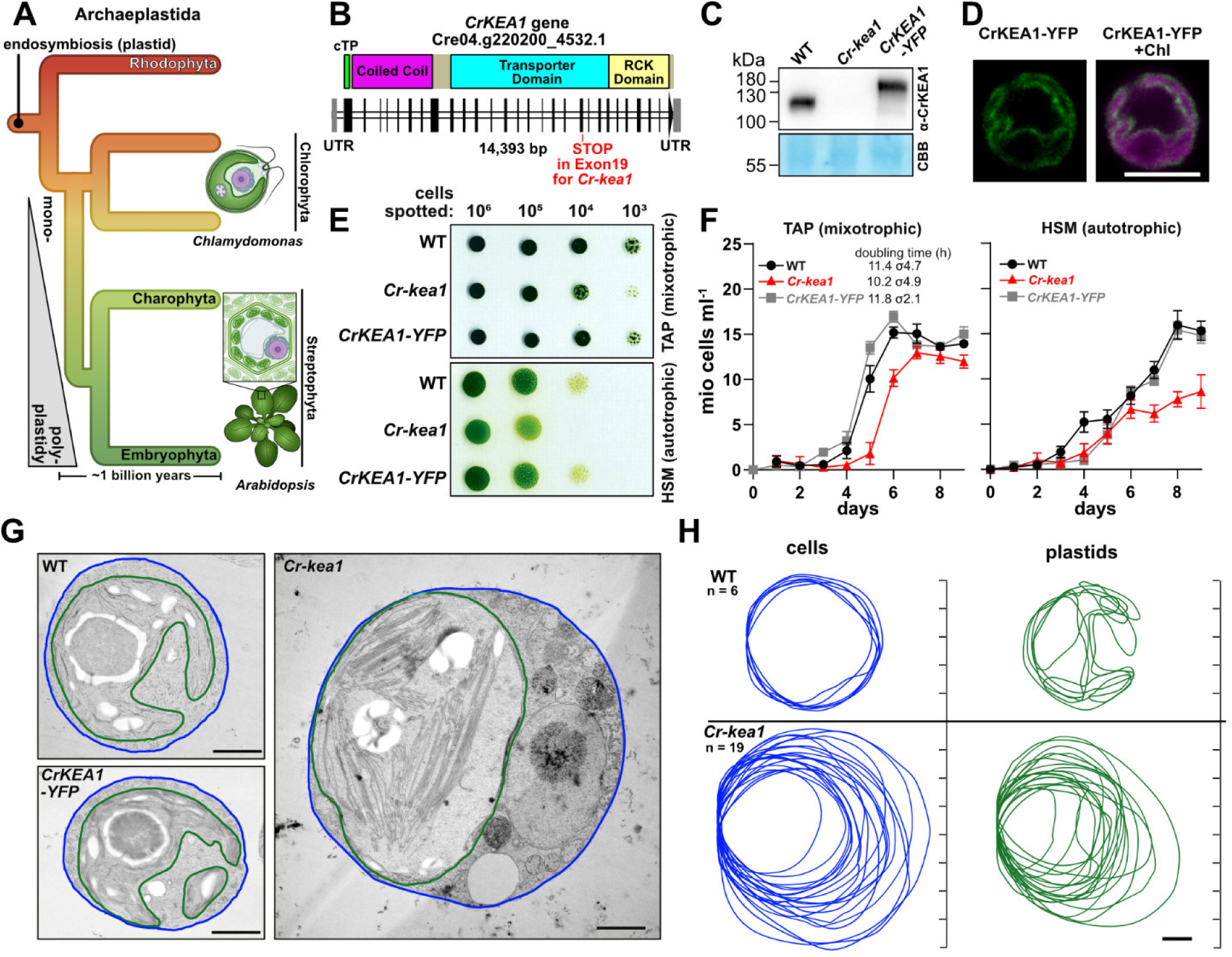
Characterization of *CrKEA1* knock-out and complementation lines: growth, localization, and cellular ultrastructure. **(A)** Simplified cladogram of the Archaeplastida showing the phylogenetic relationship between monoplastidic *Chlamydomonas* algae and multicellular, polyplastidic *Arabidopsis* plants, both derived from a common monoplastidic ancestor that diverged approximately 1 billion years ago. **(B)** Schematic of the CRISPR/Cas9 knock-out line *Cr-kea1.* **(C)** Immunoblot using anti-CrKEA1 antibodies on total protein extracts from WT, *Cr-kea1*, and the complementation line (*CrKEA1–YFP* expressed in *Cr-kea1*). **(D)** Localization of YFP-tagged CrKEA1 to the plastid envelope in the complementation line. Scale bar 5 µm. **(E)** Drop cultures grown on mixotrophic (acetate-containing TAP) and photoautotrophic (HSM) solid media at 25 µE. **(F)** Growth curves of mixotrophic and photoautotrophic liquid cultures at 25 µE. Cell counts are shown as mean ± SD (n = 3), and calculated doubling times are indicated as mean ± SD (σ). **(G)** Representative TEM images of cells grown in HSM at 100 µE. Scale bar 1 µm. **(H)** Comparison of cell and chloroplast size and shape based on TEM micrographs shown in panel G. Scale bar 1 µm.

Despite around a billion years of divergence between chlorophyte algae and streptophyte plants (Figure 1A), IE KEA transporters are conserved across the green lineage, and orthologs are present in green algae (Kunz et al., 2024). This conservation raises the question of whether KEA proteins maintain mechanistic similarity or whether their roles have been rewired in line with lineage-specific cellular contexts.

In this study, we employ Chlamydomonas for an initial investigation into how KEA-mediated ion transport across plastid envelope membranes influences the organelle’s function, cell physiology, and cell cycle in a unicellular, monoplastidic organism. By combining mutant analyses, transcriptome comparisons with Arabidopsis (DeTar et al., 2021), and cross-species complementation, we show that KEA function is a prerequisite for maintaining the integrity of ribosome biogenesis. Leveraging the complementary strengths of a multicellular polyplastidic plant and a unicellular monoplastidic alga (Gutman and Niyogi, 2004), we provide evidence that KEA’s importance for plastid gene expression is deeply conserved, while its cellular consequences are shaped by lineage-specific developmental context.

## Results

### Generation of CrKEA1 knock-out and complementation lines for subcellular localization studies of CrKEA1

*KEA* genes are conserved from green algae to angiosperms (Chanroj et al., 2012). Through amino acid sequence homology analysis, we identified Cre04.g220200 (*CrKEA1*) in Chlamydomonas as the only homolog of inner envelope (IE) K^+^/H^+^ exchangers AtKEA1/2 from Arabidopsis (see alignment, Supplemental Figure S1). The conserved monovalent cation/proton antiporter 2 (CPA2) domain and the regulatory K⁺ transport and NAD-binding (KTN) domain share 56% sequence identity and 71% similarity between plant and algae KEA1. In contrast, the extended N-terminal region (∼500 amino acids) — a characteristic feature of all IE KEAs that includes a coiled-coil domain embedded within a highly disordered segment (Chanroj et al., 2012; Bölter et al., 2020) — is far less conserved, showing only 13% identity and 29% similarity.

To investigate CrKEA1 function in Chlamydomonas, we initially generated a CrKEA1 antibody (Supplemental Figure S2, A-C) with a detection sensitivity against its antigen (CrKEA1_aa94-337) in the picogram range (Supplemental Figure S2D). We attempted to utilize mutant strain LMJ.RY0402.187220, which at the time was the only CLiP insertion line in the *CrKEA1* locus (Li et al., 2016). However, protein expression in wild-type (WT) cells and LMJ.RY0402.187220 was clearly detectable at an apparent size of ∼120 kDa (calculated size 108 kDa for full length size, without estimated chloroplast targeting peptide, cTP, ∼101 kDa), confirming functionality of the CrKEA1 antibody with Chlamydomonas protein extracts but also that the CLiP line had not lost CrKEA1 (Supplemental Figure S2E). We therefore turned towards an established CRISPR/Cas9 protocol (Kelterborn et al., 2022) to generate *CrKEA1* knock-out strains ourselves. We successfully isolated a *Cr-kea1* mutant with a stop codon insertion in exon 19 (Figure 1B), which was confirmed by PCR screening and Sanger sequencing (Supplemental Figure S3, A-D; Supplemental Table S1). Immunoblotting further verified the loss of full-length CrKEA1 (Figure 1C; Supplemental Figure S3E). Lastly, we sequenced the genome of *Cr-kea1*, which allowed us to annotate the entire insertion site (Supplemental Figure S3F, Figure S4) and to verify the absence of additional CRISPR/Cas9-mediated insertions in this background.

To analyze the subcellular localization of CrKEA1, we generated several complementation lines by transforming *Cr-kea1* with *CrKEA1* fused to a *YFP* at the C-terminus under the control of its native promoter. Immunoblotting with the CrKEA1 antibody confirmed CrKEA1-YFP expression, detecting a band at ∼140 kDa that migrated ∼30 kDa larger compared to the endogenous signal (Figure 1C; Supplemental Figure S5A) — consistent with the added YFP tag (calculated 138 kDa full length, 132 kDa without cTP). Subsequently, we performed fluorescence microscopy and found the YFP fluorescence signal to emerge from the membrane that surrounds the chlorophyll autofluorescence. This confirms that CrKEA1, consistent with its plant homologs, resides in the plastid envelope membrane (Figure 1D; Supplemental Figure S6).

### Hampered growth upon loss of CrKEA1 in Cr-kea1 strains

Chlamydomonas can grow hetero- and mixotrophically on acetate-supplemented TAP medium or autotrophically in HSM medium, relying entirely on CO₂ fixation via RuBisCO (Sager and Zalokar, 1958; Harris et al., 2009; Grossman et al., 2010; Lauersen et al., 2016). To assess the impact of CrKEA1 loss, we tested our strains under different growth conditions at 23°C. Initially, we employed agar plates and liquid cultures at 25 µmol photons m^−2^ s^−1^ (hereafter abbreviated as µE) of light to avoid stress exposure of potentially affected mutants. On TAP plates, slightly decreased growth was observed for *Cr-kea1* only at ≤ 10^4^ spotted cells. This effect was more prevalent if mutant cells were grown autotrophically on HSM plates. Here, genotype-specific differences already emerged at cell numbers between 10^6^ and 10^5^. In both cases, the expression of CrKEA1-YFP in the *Cr-kea1* background restored WT growth behavior in all four complementation lines tested (Figure 1E; Supplemental Figure S5B). The slow growth of *Cr-kea1* cells replicated well in liquid cultures. Again, the absence of acetate clearly exacerbated this effect; even after nine days of culturing, *Cr-kea1* reached only half the cell number as WT and the *CrKEA1-YFP* complemented the *Cr-kea1* strain (Figure 1F).

### Chloroplast morphology appears altered in Cr-kea1 under moderate light conditions

To investigate whether loss of CrKEA1 affects chloroplast organization, we first examined chloroplast morphology using confocal fluorescence microscopy based on chlorophyll autofluorescence (Supplemental Figure S7A, B). Under low light and photoautotrophic conditions (HSM), WT, *Cr-kea1*, and *CrKEA1–YFP* complementation lines displayed the characteristic cup-shaped chloroplast typical of Chlamydomonas. When cells were grown under moderate light, *Cr-kea1* chloroplasts appeared more heterogeneous, often with less distinct peripheral lobes but a swollen main chloroplast body compared to WT and the complemented strain.

To examine these observations at higher resolution, we performed transmission electron microscopy (TEM) on cells grown in HSM under moderate-light conditions (Figure 1G, H; Supplemental Figure S7C).

WT and CrKEA1–YFP complemented *Cr-kea1* cells consistently displayed the canonical cup-shaped chloroplast with well-defined lobes, regular thylakoid lamellae, and a distinct pyrenoid matrix. In contrast, chloroplasts in *Cr-kea1* cells appeared more spheroidal, and the characteristic lobes were not resolved in the analyzed TEM sections, although sectioning effects cannot be excluded. Thylakoid bundles within these chloroplasts also appeared more disorganized compared to the regular lamellar arrays observed in WT.

In addition, WT-like pyrenoid structures were not readily identifiable in the TEM sections of *Cr-kea1* cells. However, fluorescence imaging indicates that pyrenoid-like structures remain present in most mutant cells (detectable as gap in the autofluorescence). Finally, variation in cell and plastid size was markedly increased in the mutant population, pointing to a loss of volume homeostasis.

Together, these observations suggest that loss of CrKEA1 affects chloroplast organization and cellular morphology, particularly under photoautotrophic moderate-light conditions.

### Cr-kea1 mutants show altered K^+^ accumulation and slight cytosolic pH changes

To assess whether the growth defects of *Cr-kea1* mutants are associated with disturbed ion homeostasis, we quantified the elemental composition of whole cells grown in TAP medium at 25 µE using total reflection X-ray fluorescence (TXRF; Holzner et al., 2026).

Elemental profiles on a per-cell basis were broadly consistent with previously reported values (Merchant et al., 2020; Schmollinger et al., 2021) and showed elevated levels of several elements in the *Cr-kea1* mutant, consistent with its increased cell size (Supplemental Figure S8A). However, after normalization to cell volume, only K^+^ levels remained significantly elevated (1.85-fold) in *Cr-kea1*, whereas differences in other elements did not reach statistical significance (Figure 2A). The complementation line displayed intermediate K⁺ levels between mutant and WT. Because IE KEA transporters mediate K⁺/H⁺ exchange across the chloroplast envelope, their disruption is expected to primarily alter intracellular ion distribution rather than total cellular K⁺ content. Whole-cell measurements therefore likely underestimate the extent of ionic perturbations within subcellular compartments.

**Figure 2.**
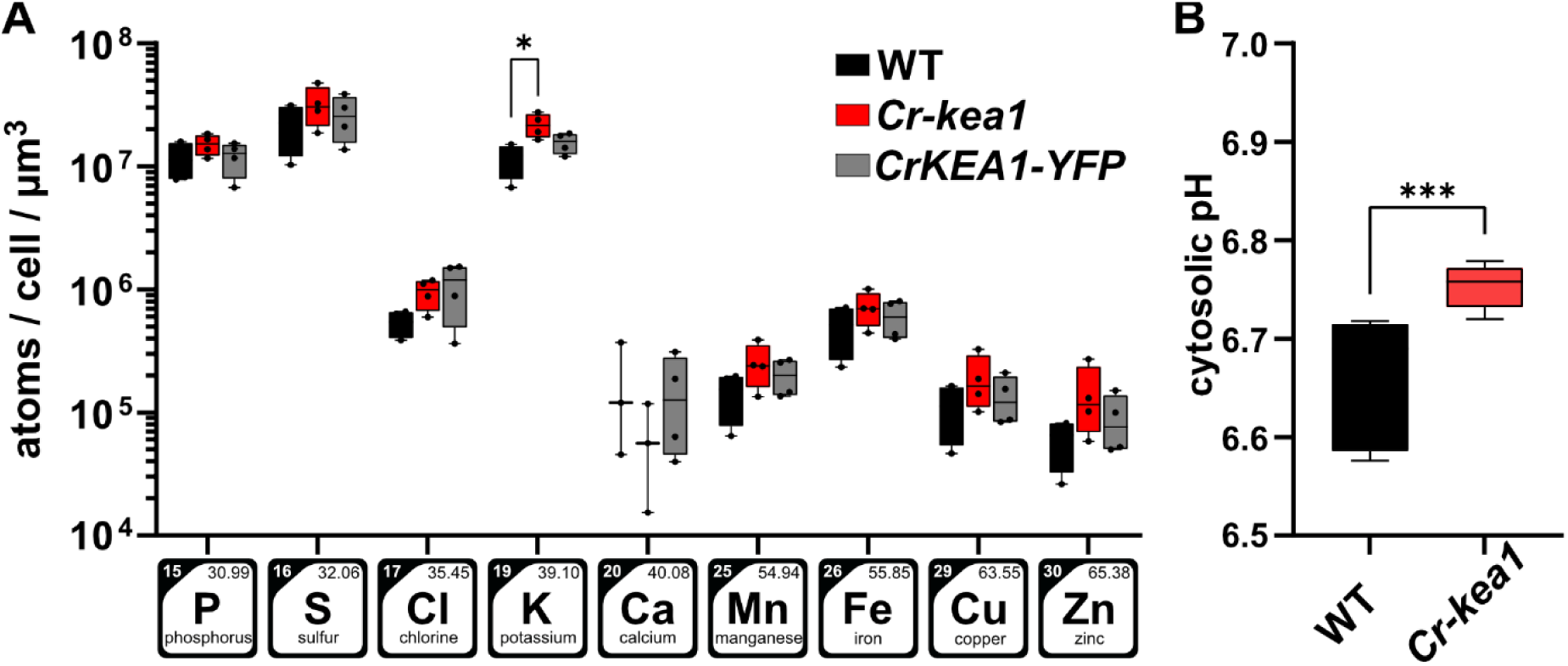
Cellular ion composition and cytosolic pH in *Cr-kea1*. **(A)** Volume-normalized ionomic profiles determined by Total Reflection X-Ray Fluorescence (TXRF). Whole-cell elemental composition was normalized to the average cell volume derived from flow cytometry measurements. Data were evaluated by one-way ANOVA for each element separately, followed by Tukey’s multiple comparison test. n = 4 **(B)** Cytosolic pH determined using the ratiometric fluorescent dye BCECF-AM. Fluorescence ratios were measured and converted to cytosolic pH values using a calibration curve generated with nigericin in buffers of defined pH. Statistical analysis was performed using unpaired t test; n = 9. Box plots show the median (center line), the 25th–75th percentiles (box), and the minimum and maximum values (whiskers). Asterisks indicate significant differences (* p < 0.05, *** p < 0.001). Cultures were grown in TAP at 25 µE prior to both measurements.

Alkali cation/H^+^ antiporters can alter pH and/or cation homeostasis (Sze and Chanroj, 2018). To determine whether the K^+^ changes in *Cr-kea1* are accompanied by altered proton homeostasis, we next measured cytosolic pH using the ratiometric fluorescent dye BCECF, which has been previously established for pH measurements in Chlamydomonas (Braun & Hegemann, 1999; Pang et al., 2023). BCECF-AM enters the cell in an esterified form and becomes trapped in the cytosol after esterase cleavage, enabling ratiometric pH determination based on the fluorescence excitation ratio (490/440 nm; Figure S8B). Using this approach, we observed a slightly elevated cytosolic pH in the *Cr-kea1* mutant compared to WT (Figure 2B). If loss of CrKEA1 leads to partial retention of K⁺ within the chloroplast, the cytosol may experience a relative K⁺ deficit and trigger increased proton extrusion via plasma membrane H⁺-ATPases to drive K⁺ uptake. This could contribute to the mildly elevated cytosolic pH observed in the *Cr-kea1* mutant.

While BCECF does not report the stromal pH, these measurements indicate that perturbation of chloroplast K⁺/H⁺ exchange in *Cr-kea1* is accompanied by measurable changes in cytosolic proton balance. Together with the altered K^+^ levels detected by TXRF, these results support the conclusion that loss of CrKEA1 leads to a broader disturbance of cellular ion homeostasis.

### Loss of CrKEA1 results in a light-dependent decrease in photosynthetic performance

The observed growth defects may correlate with impaired photosynthetic performance. Hence, we recorded electron transport rates (ETR) during step-wise light intensity (photosynthetically active radiation; PAR) increases. Interestingly, once light intensities surpassed the moderate range ≥80 PAR, *Cr-kea1* cells revealed lower ETR, consistent with increased light susceptibility compared to WT and the complementation line (Figure 3A). A similar effect was observed in drop cultures. While the maximum quantum efficiency of PSII (*F*_v_/*F*_m_) did not differ between genotypes at 0–20 µE, stepwise increases in light intensity (every hour) caused a stronger decline in *F*_v_/*F*_m_ in *Cr-kea1* compared to WT and the complementation line (Figure 3B). The same trend was observed after long-term exposure (4 days) to 25 or 100 µE in *Cr-kea1*, compared with the WT and four independent *CrKEA1-YFP* strains (Supplemental Figure S5B).

**Figure 3:**
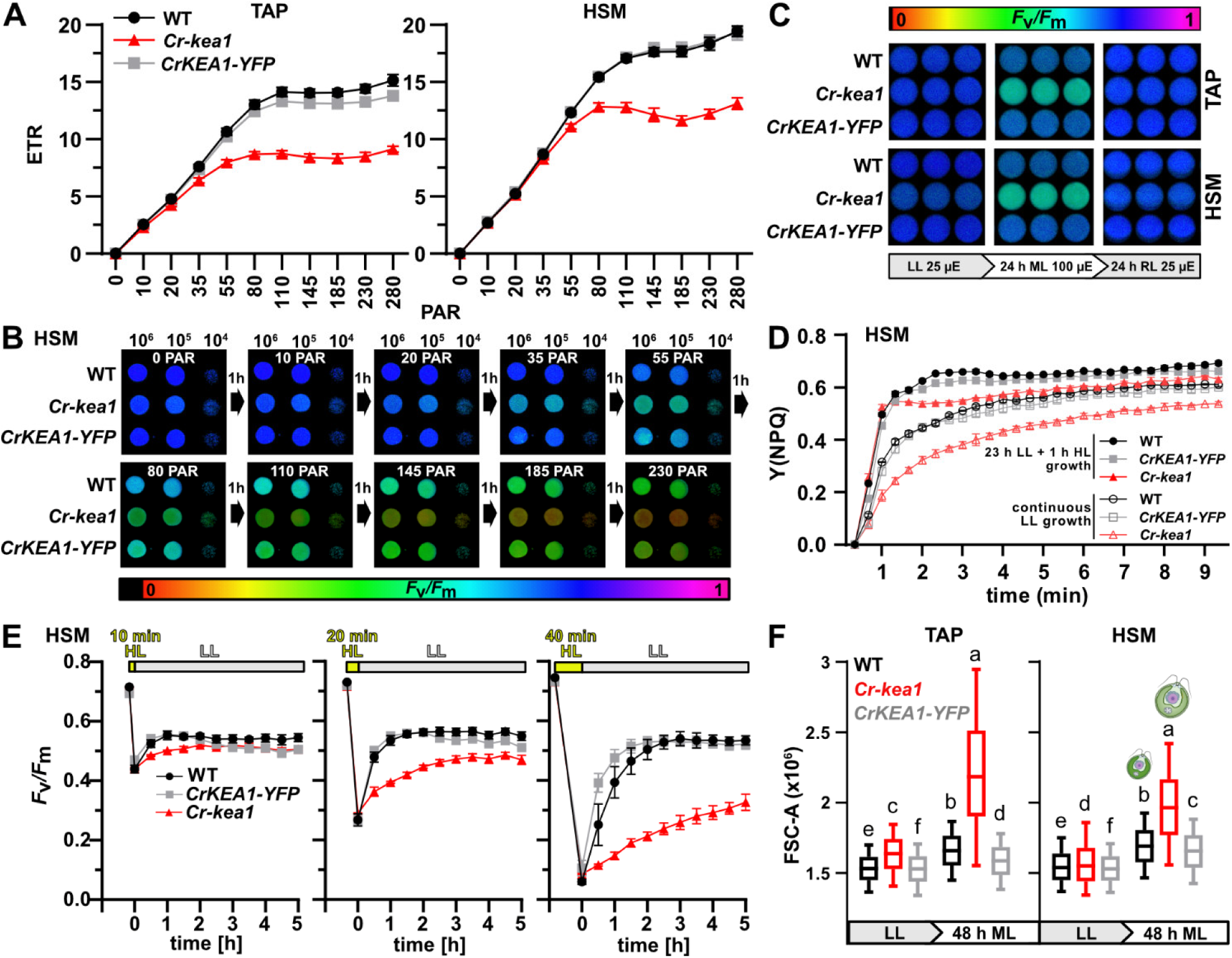
Photosynthetic characterization and cell size of *Cr-kea1*. **(A)** To assess the photosynthetic performance of *Cr-kea1*, liquid TAP and HSM cultures were exposed to gradually increasing light intensities, with PAR raised in 20-s intervals. Mean electron transfer rate (ETR) was plotted with standard deviation. n=4. **(B)** *F*_v_/*F*_m_ values were determined from HSM drop cultures every 1 h while stepwise increasing light intensity. **(C)** Light-shift experiment with TAP and HSM liquid cultures. *F*_v_/*F*_m_ was measured before and after transferring cultures from low light (LL) to moderate light (ML) for 24 h and back to low light for 24 h of recovery (RL). **(D)** Non-photochemical quenching (NPQ) induction at 646 PAR measured by PAM fluorometry. Cultures were either analyzed directly after growth in low light (LL, 25 µE) or growth under a light regime including a brief high-light (1h, 300 µE) treatment applied 15 h before measurements to induce LHCSR3 expression. Data represent mean values with standard deviation, n = 3. **(E)** Recovery from photoinhibition. Cultures were exposed to high-light treatments (646 PAR) for 10, 20, or 40 min to induce photoinhibition, followed by transfer to low light. Recovery of PSII efficiency was monitored as *F*_v_/*F*_m_ over time. Data represent mean values ± SD, n = 3. **(F)** The average cell size of *Cr-kea1* increased under moderate light (100 µE). Chlamydomonas cultures grown in HSM or TAP at 25 µE were shifted to 100 µE and analyzed by flow cytometry (FC) before and 48 h after the light shift. FSC-A values correlate with cell size. Boxes represent the interquartile range, the line indicates the median, and whiskers extend to the 10th and 90th percentiles; outliers (0–10 % and 90–100 %) are not shown. Letters indicate statistical groups. Statistical analysis was performed using Kruskal–Wallis test with Dunn’s post hoc test. Three replicates, each containing >100,000 cells, were combined for analysis and visualization. All strains were pre-cultured at 25 µE and 23 °C in TAP under shaking conditions.

For additional insight, we established a light intensity shift routine for liquid cultures. Initially, *F*_v_/*F*_m_ was determined in all three genotypes grown at 25 µE either in HSM or TAP media. Regardless of the growth media used, no clear differences emerged across the strains. However, if the same genotypes were shifted to standard moderate light conditions (100 µE; Bialevich et al., 2022) for 24 h, a clear *F*_v_/*F*_m_ drop was noticeable for the mutant but not for the WT and all six *CrKEA1-YFP* lines measured (Figure 3C; Supplemental Figure S5C). Subsequent incubation for 24 h at 25 µE restored *F*_v_/*F*_m_ to baseline levels.

To further investigate the physiological basis of the reduced ETR and *F*_v_/*F*_m_ values observed under moderate light, we assessed non-photochemical quenching (NPQ) induction and recovery from photoinhibition. Because NPQ capacity in Chlamydomonas depends on the high-light–inducible LHCSR3 protein (Peers et al., 2009), NPQ was measured both in low-light–grown cultures and in cultures subjected to repeated brief high-light treatments to induce LHCSR3 expression. Under both conditions, *Cr-kea1* cells displayed reduced NPQ induction compared to WT and the complementation line, although NPQ increased in all strains following high-light pretreatment (Figure 3D).

We next examined recovery from photoinhibition by exposing cultures to high-light treatments that caused a defined decrease in *F*_v_/*F*_m_, followed by transfer to low light while monitoring PSII recovery over time. In these experiments, WT and the complementation line recovered more rapidly than the *Cr-kea1* mutant (Figure 3E).

In summary, *Cr-kea1* mutants are highly light sensitive and show clear photosynthetic defects, which are fully rescued by *CrKEA1-YFP* expression, confirming loss of CrKEA1 as the causative mutation. While low light intensities (25 µE) are mostly permissive for *Cr-kea1*, PSII efficiency declines sharply under moderate light (∼100 µE). Consistent with this, *Cr-kea1* cells display reduced NPQ capacity and recover more slowly from photoinhibition, indicating that the mutant is less effectively protected from light-induced damage and has a reduced capacity to restore PSII activity after stress.

### Flow cytometry shows increased cell sizes for Cr-kea1 shifted from low to moderate light

Next, we examined the response of TAP and HSM cultures to the light shift using flow cytometry (Figure 3F; Supplemental Figure S9). Regardless of the growth medium, *Cr-kea1* cells were slightly larger in size based on FSC-A values at low light (25 µE). 48 h after the shift to moderate light (100 µE), cell size increased for both media across all genotypes. However, *Cr-kea1* cultures showed the strongest increase in cell size and broadening of the size distribution. This phenotype was rescued in all six complementation lines examined (Supplemental Figure S5C). After 48 h of moderate light treatment, calibration indicated average volume increases of 19% in TAP and 23% in HSM for WT, 9% in TAP and 19% in HSM for the *CrKEA1-YFP* line, and 97% in TAP and 67% in HSM for *Cr-kea1* (Supplemental Table S2). Using SSC-A as a proxy for cell complexity, changes in the *Cr-kea1* mutant (82%) by far exceeded those in WT (-4%) and the complemented line (6%) under TAP conditions. A similar trend was observed in HSM, with complexity increases of 52% for *Cr-kea1*, 19% for WT, and 4% for the *CrKEA1-YFP* line (Supplemental Table 3).

### Transcriptome profiling during light acclimation reveals that the absence of CrKEA1 alters ribosome biogenesis, plastid fission, and cell cycle regulation

Given the impaired physiological performance of *Cr-kea1* mutant cells upon shift into moderate light, we sought to identify biological processes affected by the loss of *CrKEA1* through characterizing the underlying transcriptional responses. To this end, we performed RNA sequencing (RNA-seq) on cultures subjected to the established light shift regime. Cells grown under low light (LL, 25 µE) were exposed to moderate light (ML, µE) for 8 h, followed by a 24 h low-light recovery phase (RL, 25 µE) (Figure 4A). The experiment was set up with TAP medium to ensure that *Cr-kea1* cells experience stress but not the exacerbated defects observed in HSM. Ribo-depleted libraries were prepared to capture the full extent of transcriptional changes, including organellar transcripts.

**Figure 4:**
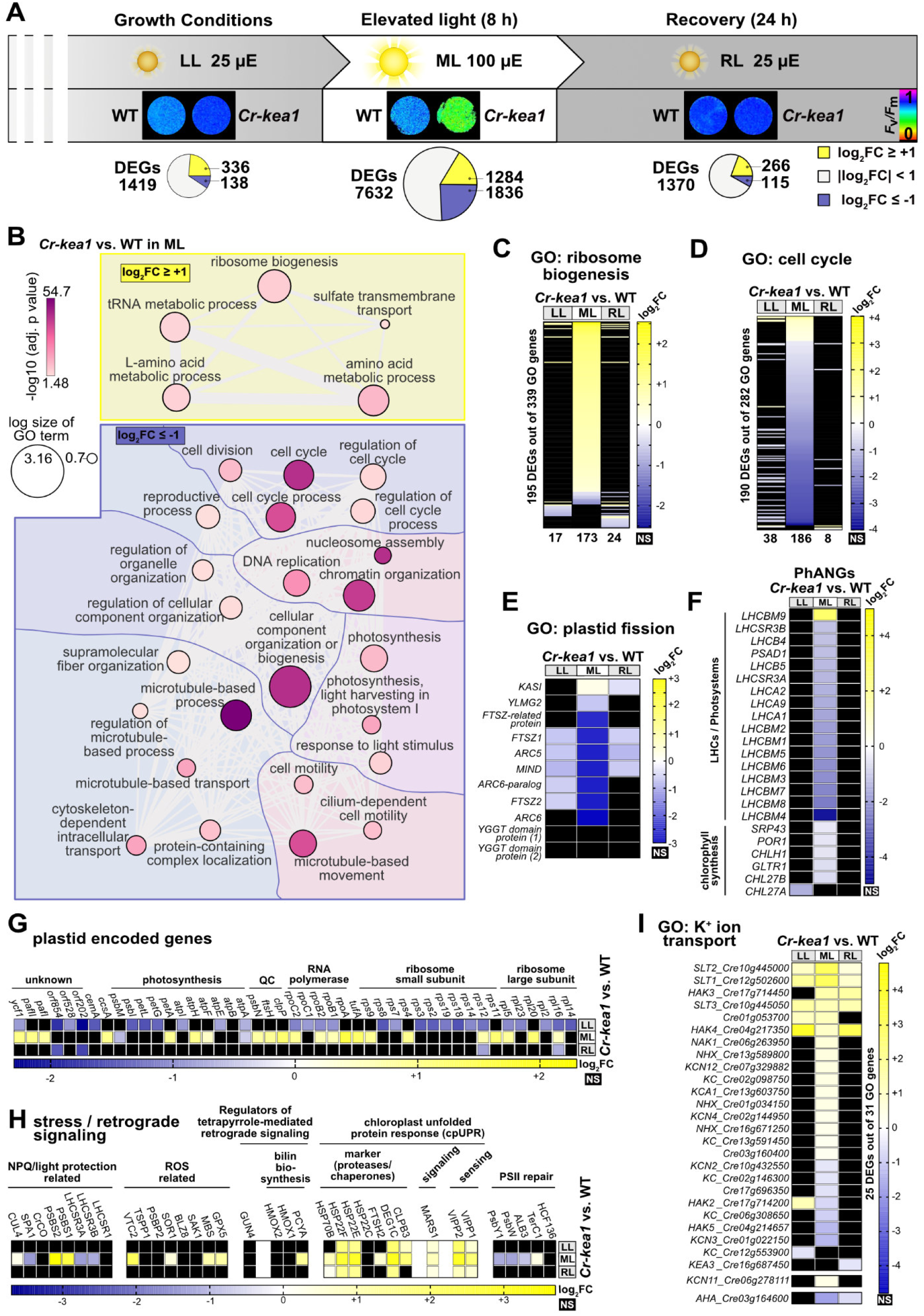
RNA-seq analysis of a light-shift experiment comparing transcriptional responses of *Cr-kea1* and WT. **(A)** Experimental setup for the RNA-seq analysis, including *F*_v_/*F*_m_ values of the cultures and pie charts indicating the number of significantly differentially expressed genes (DEGs; adj. p-value < 0.05) per steady-state comparison (*Cr-kea1* vs. WT). Yellow and blue sectors denote up- and downregulated genes with |log₂FC| ≥ 1, respectively. n = 4. **(B)** Gene Ontology (GO) terms of the *Biological Process* domain enriched among DEGs as interactive graph (REVIGO). **(C–I)** Heatmaps showing log₂FC values (*Cr-kea1* vs. WT) of significant DEGs (adj. p-value < 0.05) associated with selected GO terms or gene lists. Transcripts not reaching statistical significance in a given condition are shown in black (not significant, NS). “GO genes” indicates the number of genes associated with each respective GO term. (C) Ribosome biogenesis (GO:0042254). (D) Cell cycle (GO:0007049). (E) Plastid fission (GO:0043572). (F) Photosynthesis-associated nuclear genes (PhANGs). (G) Plastid-encoded genes. (H) Selected transcripts related to stress/retrograde signaling. (I) Ion transport (including GO:0006813, potassium ion transport). NHX, sodium–hydrogen exchangers; SLT, sodium/sulfate co-transporters; KC(N), K^+^ channels; HAK, high-affinity K^+^ transporters; NAK, Na^+^– K^+^ pumps; AHA, plasma membrane proton pumps. Gene IDs and descriptions corresponding to the gene names displayed are listed in the Supplemental Table S6.

For physiological validation of the light treatment, we again recorded *F*_v_/*F*_m_ at each sampling point. While there was no genotype-specific difference at the onset of the experiments i.e., at 25 µE, only 8 h at 100 µE induced clear PSII photoinhibition (*F*_v_/*F*_m_ drop) in *Cr-kea1* mutants. By contrast, WT cells readily acclimated to the changing light regime (Figure 4A; Supplemental Figure S10A). Following the RL phase, PSII quantum yield was fully restored in all genotypes.

Multidimensional scaling (MDS) showed tight clustering of replicates (*n*=4) and a stronger genotype separation under ML (Supplemental Figure S10B). Consistent with this, the number of significantly differentially expressed genes (DEGs, adjusted p-value < 0.05) under LL (1419 DEGs) went up >6-fold under ML (7632 DEGs) but back down at RL conditions (1370 DEGs) (Figure 4A; Supplemental Figure S10C, D; Supplemental Table S4). In addition, ML DEGs were more strongly deregulated i.e., more DEGs above the |log_2_FC| ≥ 1 threshold than under other light conditions.

Gene Ontology (GO) enrichment analyses highlighted a coordinated shift in cellular programs under ML (Figure 4B; Supplemental Table S5): Upregulated DEGs were strongly enriched for *ribosome biogenesis*, including *rRNA processing*. Both categories contained cytosolic and plastid ribosome biogenesis components. Downregulated categories included *cell cycle*, *DNA replication*, *chromatin organization*, and *microtubule-based processes*, which are partly intertwined.

Consistent with this, these trends were confirmed by a detailed analysis of transcript changes. Supplemental Table S6 provides Gene IDs and annotations for key genes highlighted in the text and figures. For the heatmaps shown in Figures 4C and 4I, again only significantly differentially expressed genes (adjusted p value < 0.05) were considered for interpretation; genes that did not meet this threshold in a given condition are indicated in black. Under these conditions, a broad upregulation of ribosome-related genes was observed when *Cr-kea1* cells were exposed to moderate light (Figure 4C). By contrast, cell cycle–related genes were by and large downregulated in *Cr-kea1* cells under these light conditions (Figure 4D). While *CDKA1*, the major driver of the G1/S transition (Tulin and Cross, 2014), was unchanged, multiple key regulators of the Chlamydomonas cell cycle were expressed below WT-level in *Cr-kea1*: Expression of genes controlling S/M transitions by blocking spindle formation was strongly decreased, including *CDKB1(2)* and its activating cyclins (*CYCA1*, *CYCB1*). Three of the D-type cyclins, which mediate G1/S entry (Bisova et al., 2005), were mildly reduced (∼2-fold for *CYCD3, 5*). In addition, transcript accumulation for both *CDKG1*, which becomes induced when cells reach sufficient size for division (Li et al., 2016), and for *MAT3*, a retinoblastoma homolog regulating the commitment to division (Umen and Goodenough, 2001), were slightly diminished. Together, these changes provide evidence that the cell cycle entry and S/M cycling might be affected in the *Cr-kea1* mutants (Supplemental Figure S10E).

Interestingly, most transcripts associated with the GO term *plastid fission* were markedly reduced in light-shifted *Cr-kea1* cells (Figure 4E), suggesting a link between KEA1 function in the plastid envelope membrane and chloroplast division control in algae.

### Absence of CrKEA1 suppresses PhANGs but triggers an induction of PGE and protein homeostasis genes

Arabidopsis *kea1kea2* mutants display delayed chloroplast biogenesis caused by activation of GUN1-dependent retrograde signaling, which suppresses the expression of GOLDEN2-LIKE transcription factors and their downstream gene cluster of photosynthesis-associated nuclear genes (PhANGs; DeTar et al., 2021). Hence, we curated a *PhANG* list for Chlamydomonas. Interestingly, *Cr-kea1* cells showed a clear repression of genes encoding for light-harvesting complexes, photosystem subunits, and chlorophyll biosynthesis enzymes under ML (Figure 4F), which is reminiscent of the response described for *At-kea1kea2* plant mutants (DeTar et al., 2021). Furthermore, among downregulated genes, also photosynthesis-related GO terms were enriched in the *Cr-kea1* mutant (Figure 4B). By contrast, plastid-encoded photosynthesis genes were largely unaffected, though transcripts for plastid transcription/translation components (e.g., RNA polymerase subunits, ribosomal proteins), essential for plastid gene expression (PGE), were elevated in *Cr-kea1* compared to WT (Figure 4G; Supplemental Figure S10G). This suggests that the photosynthetic decline may not only arise from a transcriptional repression within the plastid, but in addition from defective translation, consistent with the general enrichment of ribosome biogenesis transcripts.

Other DEG categories were more selective. Chloroplast-encoded genes for chlorophyll biosynthesis were unchanged in *Cr-kea1* (Supplemental Figure S10G). Likewise, *HMOX* and *GUN4*, which regulate tetrapyrrole and bilin synthesis relevant for retrograde signaling in Chlamydomonas (Duanmu et al., 2013), were unaffected. However, bilin-synthesis gene *PCYA* was upregulated under ML (Figure 4H). Chlamydomonas lacks close *GOLDEN2-LIKE transcription factor* homologs and probably relies mainly on a bilin-derived retrograde signaling pathway to repress PhANGs and facilitate acclimation (Duanmu et al., 2013; Richter et al., 2023). Hence, the observed *PCYA* induction likely contributes to the *PhANG* suppression observed in *Cr-kea1*.

Next, we examined nuclear genes implicated in protein homeostasis, ROS defense, and photoprotection (Figure 4H). Plastid-encoded quality control proteins, namely *PsbN* involved in PSII repair and the proteases *ClpP* and *FtsH*, were mildly upregulated in *Cr-kea1* under ML, while nuclear-encoded PSII repair factors (*HCF136*, *TerC1*, *ALB3*, *PsbW*, *PsbY*; Plöchinger 2016) showed no consistent change with a tendency for down-regulation. Several ROS-related genes (*GPX5*, *MBS*, *SOR1*, *VTC2*) were upregulated in the mutant under ML, whereas photoprotection factors (qE/NPQ) showed mixed patterns: *PSBS* transcripts were elevated in the mutant, but *LHCSR* induction was attenuated relative to WT, consistent with the reduced NPQ observed in *Cr-kea1* (Figure 3D). In turn, *SPA1*, a component of the E3 ubiquitin ligase complex (CUL4-DDB1-LRS1-SPA) involved in repression of PSBS- and LHCSR-mediated qE (Gabilly et al., 2019; Aihara et al., 2019), was downregulated, implying altered control of qE responses in *Cr-kea1*.

Because higher expression of protein quality control genes (*ClpP* and *FTSH*) was observed in the plastid, we also analyzed markers of the chloroplast unfolded protein response (cpUPR). Genes encoding cpUPR components, including *VIPP2* (sensing), *MARS1* (signaling), and markers such as *CLPB3*, *DEG1C*, *FTSH2*, *HSP22E/F*, and *HSP70B* (Ramundo et al., 2014; Perlaza et al., 2019; Theis et al., 2020), were consistently upregulated in *Cr-kea1* compared to WT (Figure 4H), in most cases already at LL conditions i.e., 25 µE.

### Expression of ion transporters in Cr-kea1 is altered under low and moderate light

When analyzing genes upregulated (log₂FC ≥ 1) in *Cr-kea1* LL conditions, the GO term potassium transport (GO:0006813) was significantly enriched (Supplemental Table S5). Within this group, several HAK/KUP/KT-type high-affinity K⁺ transporters (*HAK2*, *HAK4*) and Na⁺/SO₄²⁻ transporter-like genes (*SLT1-SLT3*) showed elevated transcript levels in the mutant compared to WT (Figure 4I, Supplemental Figure S10F). These transporter families have previously been linked to K⁺ and sulfate acquisition and to the maintenance of cellular ion balance (Li et al., 2023; Pootakham et al., 2010).

Inspection of the complete gene set within the K^+^-transport category revealed that expression changes persisted or became more pronounced under moderate light (ML). The same transporters that were induced under LL remained among the most strongly upregulated genes under ML, except *HAK2*, which decreased relative to WT. Additional transporters, including *HAK3* and *NAK1*, the sole Na⁺/K⁺-ATPase-like pump homolog in Chlamydomonas, were also induced. NAK1 and HAK-like proteins respond to low- K^+^ and saline stress and mediate cytosolic K⁺ and Na⁺ homeostasis (Li et al., 2023). Hence, their upregulation in *Cr-kea1* suggests a disturbance in cellular ion homeostasis and activation of low-K^+^–responsive pathways, possibly reflecting altered K⁺ partitioning causing cytosolic K^+^ deficiency or impaired ion signaling between the chloroplast and cytosol (Figure 4I).

Across the light-shift experiment, *SLT* and *HAK* family members, along with other significant genes from this GO term, followed a similar trend in *Cr-kea1* (and partially in WT) in response to the light treatment: transcript levels increased under ML and declined again during recovery (RL), but transcription was constantly higher in the *Cr-kea1* mutant than in the WT (Supplemental Figure S10F).

*KCN11*, a tonoplast K⁺ channel required for osmoregulation, also showed significantly elevated transcript levels (log_2_FC = 0.43) in *Cr-kea1* under ML (Figure 4I). The upregulation of KCN11 in *Cr-kea1* thus supports the interpretation that loss of *CrKEA1* perturbs osmotic balance, consistent with broader alterations in ion homeostasis.

In summary, loss of *CrKEA1* triggers a multifaceted transcriptional response pointing towards a dual role for CrKEA1-dependent ion homeostasis in maintaining chloroplast translation capacity and safeguarding protein folding.

### Species-Specific Consequences of CrKEA1 Deficiency for Plastid and Cell Cycle Regulation

Having observed similar transcriptional upregulation of ribosome biogenesis genes in both species upon *KEA* loss, we aimed to explore similarities and differences in transcriptomic responses between Chlamydomonas and Arabidopsis IE *KEA* knockouts. To this end, we re-analyzed RNA-seq data from Arabidopsis WT and *At-kea1kea2* mutants (DeTar et al., 2021; Supplemental Table S7) and compared it with the RNA-seq data from the Chlamydomonas *Cr-kea1* mutant. To identify genes commonly or uniquely regulated in response to IE *KEA* disruption across species, we mapped Chlamydomonas DEGs to their closest Arabidopsis homologs and categorized the results into three groups: shared DEGs (genes differentially expressed in both species) and species-specific DEGs (Supplemental Table S8).

Notably, transcriptomic comparisons between Arabidopsis and Chlamydomonas are inherently constrained by major biological and experimental differences between the two systems, including fundamental differences in physiology, cellular organization, and organism-specific growth conditions (Julca et al., 2023). The cross-species comparison should therefore be interpreted as an exploratory analysis aimed at identifying broadly conserved or lineage-specific responses to IE KEA loss rather than as a strict one-to-one comparison of transcriptional regulation. Nonetheless, similarities observed despite these differences may highlight robust aspects of the cellular response to disrupted chloroplast ion homeostasis.

GO term enrichment analyses on these categories (Figure 5A; Supplemental Table S9) revealed that DEGs involved in ribosome biogenesis were largely shared between the two species, indicating a conserved stress response linked to impaired rRNA maturation. In contrast, DEGs related to the cell cycle were exclusively found in Chlamydomonas, suggesting a lineage-specific consequence of *CrKEA1* loss on cell division control.

**Figure 5:**
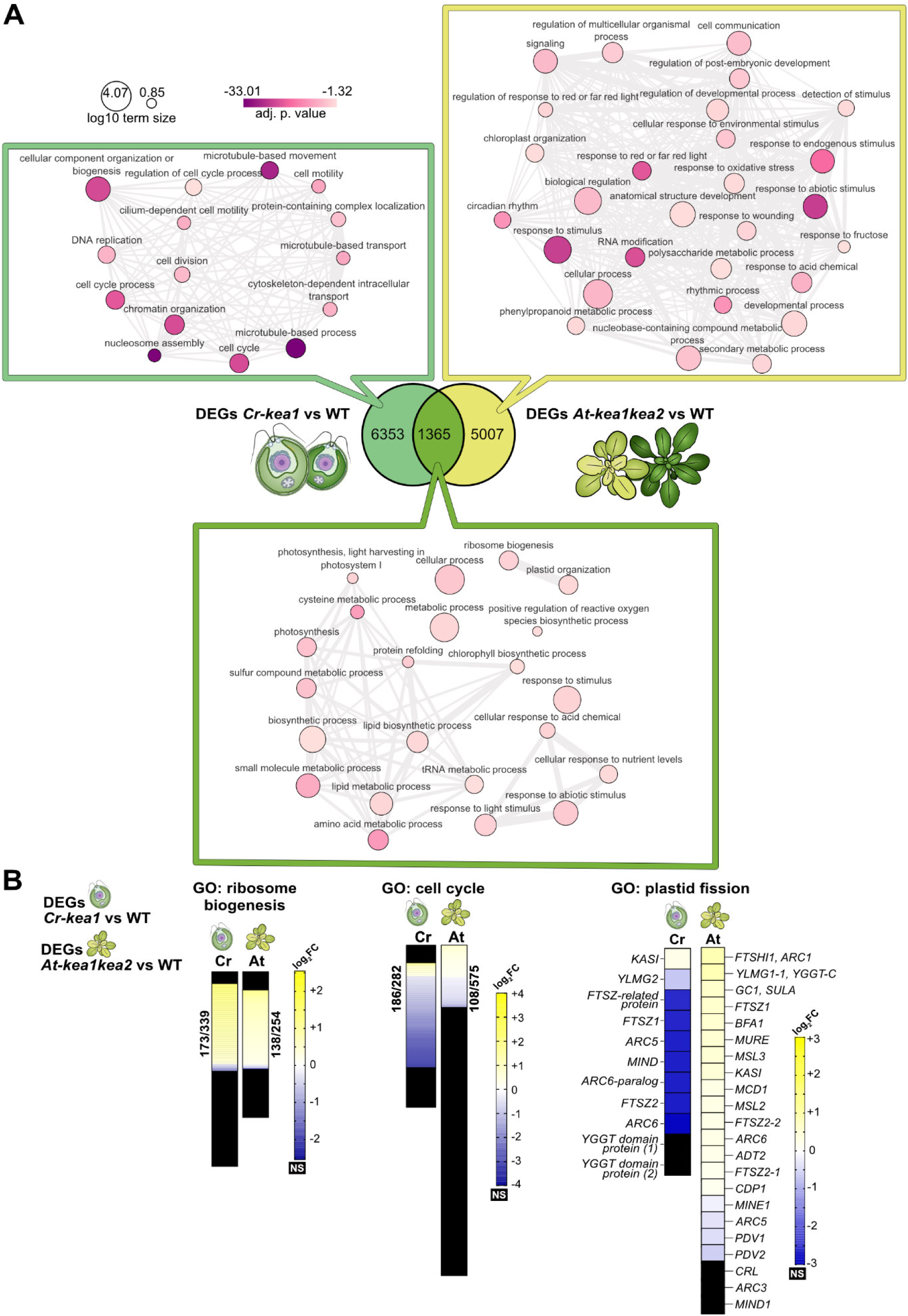
Cross-species transcriptomic comparison of inner envelope *KEA* mutants in Chlamydomonas and Arabidopsis. **(A)** Mutants lacking inner envelope *KEA* transporters (Chlamydomonas *Cr-kea1* and Arabidopsis *At-kea1kea2*) in both Chlamydomonas and Arabidopsis share a subset of DEGs. Shared DEGs were identified by mapping Chlamydomonas genes to Arabidopsis orthologs using g:Orth. Significant DEGs (adj. p-value < 0.05; numbers shown in the Venn diagram) were filtered using |log₂FC| ≥ 0.75 and subjected to Gene Ontology (GO) enrichment analysis with g:Profiler (*Biological Process* domain). GO terms are shown as interactive graph (REVIGO). Chlamydomonas DEGs correspond to the moderate light (ML) condition. Shared DEGs are enriched for *ribosome biogenesis* (GO:0042254), whereas enrichment for *cell cycle* (GO:0007049) is observed only in Chlamydomonas. **(B)** Heatmaps display log₂FC values for all genes annotated to *ribosome biogenesis*. Only significant DEGs (adj. p-value < 0.05) are color-coded, while non-significant (NS) or undetected genes are shown in black. Numbers indicate the count of significant DEGs relative to the total number of genes annotated to the respective GO term (DEGs / GO term size).

The Arabidopsis-specific DEGs were enriched for GO terms associated with, i.e., responses to various stimuli, including light, red and far-red light signaling, wounding, and regulation of multicellular development, consistent with the plants’ complex tissue architecture and photomorphogenic signaling networks (Figure 5A).

A direct comparison at the level of DEGs further confirmed that ribosome biogenesis genes are upregulated in both species (Figure 5B). However, cell cycle–related genes are strongly downregulated in Chlamydomonas but not significantly altered in Arabidopsis. Interestingly, most genes related to plastid fission showed opposite trends: they were downregulated (*MINE*, *MIND*, *FTSZ1*, *ARC5*, *ARC6*) in Chlamydomonas but many slightly upregulated in Arabidopsis (*FTSZ1*, *ARC6*, which are conserved in both) or not-significant (*MIND*). However, key plastid division genes, like *PDV1* and *PDV2*, which are unique to land plants and determine plastid division rate (Miyagishima et al., 2006; Okazaki et al., 2009, Holtsmark et al., 2013), and *MINE* and *ARC5*, were slightly down-regulated in Arabidopsis.

### Plastid rRNA processing is impaired in both Arabidopsis and Chlamydomonas IE KEA-deficient mutants

Given the transcriptional upregulation of ribosome biogenesis genes in both Arabidopsis and Chlamydomonas IE *KEA* mutants (Figure 5B), we examined whether loss of *CrKEA1* in algae also disrupts plastid rRNA maturation, leading to the accumulation of unprocessed rRNA precursors as observed in plants (DeTar et al., 2021; DeTar et al., 2022). Notably, plastid rRNA processing differs between the two lineages in several important aspects: 1) Arabidopsis generates a 4.5S rRNA fragment from the 23S precursor, whereas Chlamydomonas instead produces the distinct 7S and 3S fragments cleaved from the opposite end (Holloway and Herrin, 1998; Ahmed et al., 2016; Schmid et al., 2024); 2) Chlorophytes employ much more octotricopeptide repeat (OPR) RNA-binding proteins (RBP) while plants rely heavily on pentatricopeptide repeat (PPR) proteins (Small et al., 2023). However, both systems rely on precise cleavage of large precursor transcripts into functional ribosomal RNAs, which involves temporal sequence-specific RNA- RBP interactions (Supplemental Figure S11).

To test for stromal rRNA maturation defects in Chlamydomonas, we analyzed total RNA from *Cr-kea1* and WT cultures using rRNA sequencing. Initially, all stromal rRNA reads from *Cr-kea1* and WT were aligned akin to previous studies (Hotto et al., 2015; Castandet et al., 2016). This approach revealed increased read coverage within intergenic regions in *Cr-kea1*, particularly in the 7S and 3S regions derived from the 23S precursor. Since these fragments are removed during rRNA maturation, their high abundance in *Cr-kea1* indicated mutant-specific accumulation of unprocessed rRNA intermediates (Figure 6A, Supplemental Figure S12A, B). To corroborate these observations, we carried out RNA blots employing stromal rRNA-specific probes. Indeed, a clear accumulation of the 7S-3S-23S precursor was specifically observed in *Cr-kea1* cells (Figure 6B, Supplemental Figure S13A, B). Splicing defects did not change with light shifts, but they were less abundant in cells grown in TAP medium.

**Figure 6:**
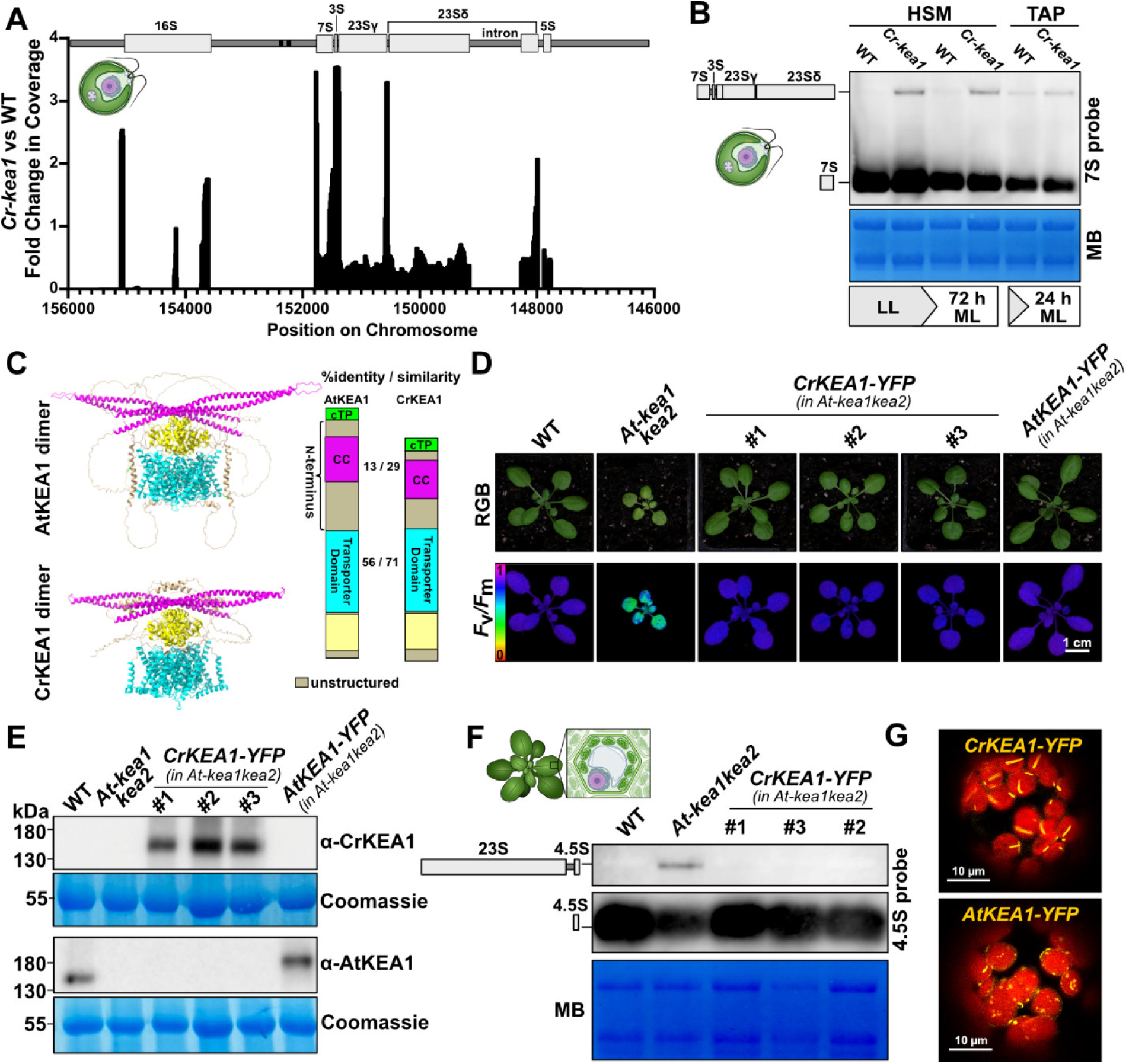
Conserved role of inner envelope KEA transporters in plastid rRNA processing across algae and higher plants. **(A)** Impaired plastid rRNA processing in *Cr-kea1* leads to enrichment of the 7S–3S–23S precursor. Shown is the fold change in normalized read coverage (*BamCompare*) from an rRNA-seq experiment comparing *Cr-kea1* and WT, revealing increased read density over intergenic regions corresponding to the 7S–3S–23S precursor in the mutant. **(B)** RNA blot analysis using specific plastid rRNA probes confirms accumulation of the 7S–3S–23S precursor in *Cr-kea1*. MB, Methylene Blue. **(C)** Structural comparison of AtKEA1 and CrKEA1. AlphaFold3 models of dimeric AtKEA1 and its algal homolog CrKEA1 are shown, with colors corresponding to domain organization depicted to the right. The representation as a dimer was chosen due to the structural findings for the bacterial homolog KefC (Gulati et al., 2024). **(D)** Phenotypical complementation of the *At-kea1kea2* mutant by expressing *CrKEA1–YFP*. Scale bar 1 cm. (**E)** Immunoblot validation of *CrKEA1–YFP* expression in complementation lines. (**F)** RNA blot demonstrating restoration of normal plastid rRNA processing in *At-kea1kea2* upon expression of *CrKEA1–YFP*. MB, Methylene Blue. (**G)** Confocal microscopy of protoplasts isolated from *At-kea1kea2* plants expressing either *AtKEA1–YFP* or *CrKEA1–YFP*. Chlorophyll autofluorescence (*red*) and YFP signal (*yellow*) displayed in false colors. Scale bar 10 µm.

In summary, results from our RNA analyses suggest that despite significant differences in rRNA structures and processing pathways between algae and land plants, the importance of IE KEAs for seamless plastid rRNA processing is functionally conserved between chlorophytes and streptophytes.

### Functional conservation of IE KEA transporters between algae and higher plants

The rRNA processing phenotypes observed in plants and green algae suggest functional conservation of IE KEAs across the green lineage. We therefore tested whether CrKEA1 can substitute for its homologs AtKEA1/2 in Arabidopsis. Despite substantial amino acid sequence divergence between CrKEA1 and AtKEA1, structural predictions generated with AlphaFold3 (DeepMind; Abramson et al., 2024) indicate a broadly conserved domain architecture (Figure 6C; Supplemental Figure S14A-C; Gulati et al., 2024). Initially, we cloned full-length *CrKEA1* with a C-terminal YFP. The finished construct *CrKEA1-YFP* was transformed into the *At-kea1-1kea2-1* (hereafter simply *At-kea1kea2*) Arabidopsis mutant allele (Kunz et al., 2014). Several independent transgenic *CrKEA1-YFP* lines exhibited robust complementation i.e., growth, leaf area, fresh weight, and photosynthetic performance (*F*_v_/*F*_m_ and ETR) were restored close to WT levels (Figure 6D; Supplemental Figure S15). Subsequent immunoblotting confirmed the accumulation of CrKEA1-YFP in the transgenic lines at the expected apparent size of ∼140 kDa (Figure 6E).

Next, we conducted RNA blots with WT, *At-kea1kea2*, and *CrKEA1-YFP* complemented lines and found that CrKEA1-YFP restores plastid 23S rRNA processing (Figure 6F).

Together, these findings demonstrate that despite a billion years of divergence, CrKEA1 retains sufficient functional similarity to AtKEA1/2, which reveals a strong evolutionary conservation of K^+^/H^+^-mediated effects on ribosome biogenesis. Curiously, when expressed in Arabidopsis, CrKEA1-YFP localized in discrete, rod-like structures, whereas the endogenous AtKEA1-YFP appeared in discrete spots (Figure 6G).

### Comparison of co-expression networks reveals a divergent cell cycle embedding in Arabidopsis and Chlamydomonas

Since the GO terms *cell cycle* and *plastid fission* were affected in the Chlamydomonas mutant but not in Arabidopsis, we carried out an in-depth comparison between the respective co-expression networks (Figure 7A; Supplemental Tables S6 and S10). Direct co-expression involves “*gene to gene*” interactions, while secondary co-expression involves “*gene to intermediary gene to gene*” interactions. In the Chlamydomonas co-expression network (ML condition based), *ARC5*, *ARC6*, and *MIND,* all scoring a log_2_FC < -2, were found to be secondary neighbors in a shared cluster with cell cycle genes: e.g., *MIND* with the key *cell cycle* gene *CDKB1(2)* but also *ARC5* with *TOPII*, *TPX2*, *AUR2*, *MCM5*, *WEE1* that were similarly downregulated with a log_2_FC < -1. This indicates that both biological processes have tightly co-expressed interactions resulting in unidirectional expression changes.

**Figure 7:**
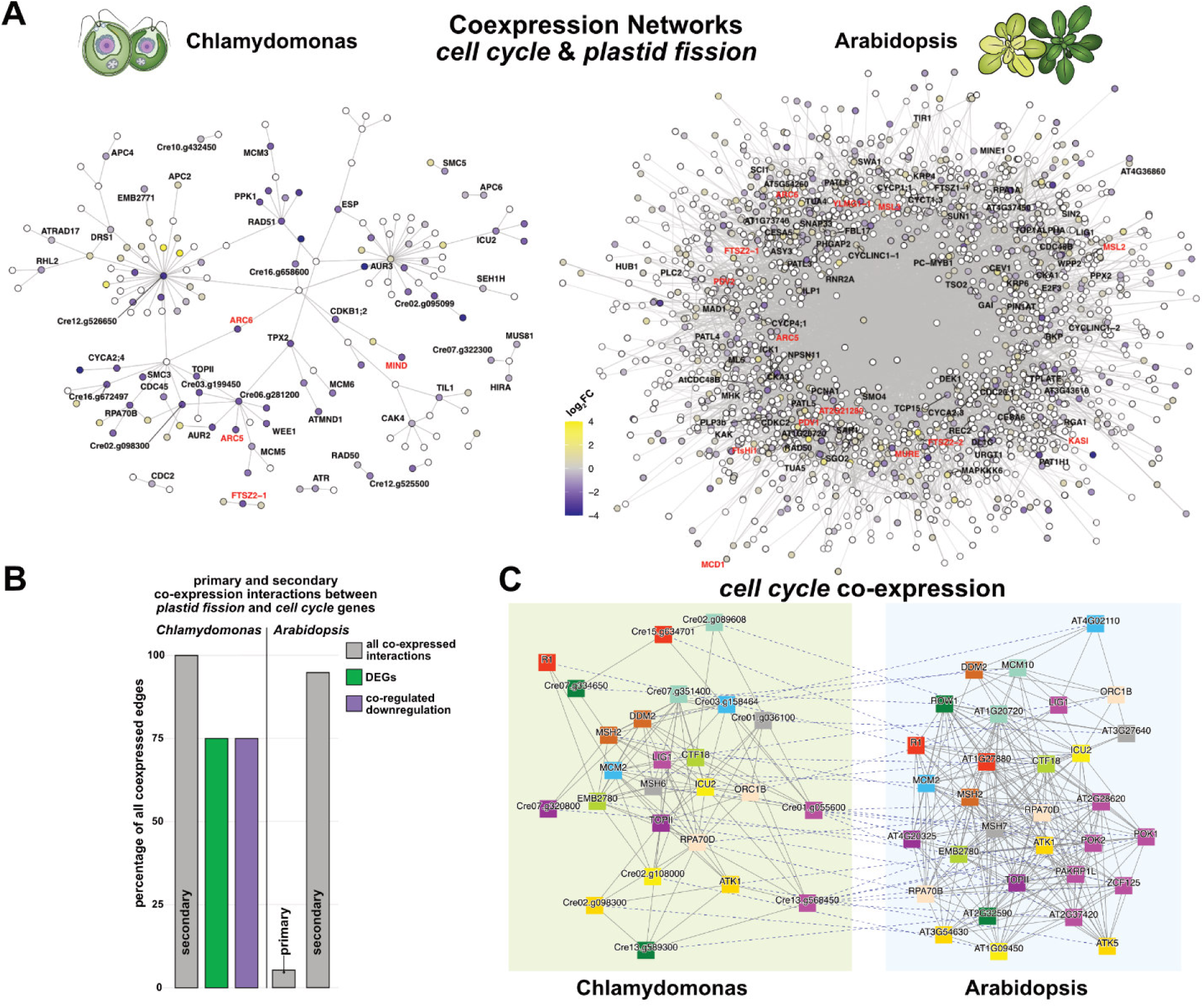
Comparisons between co-expression networks of Arabidopsis and Chlamydomonas. **(A)** From both species, genes annotated to GO:0007049 (*cell cycle*) and GO:0043572 (*plastid fission*) were extracted and visualized based on co-expression relationships in each species. Nodes represent genes and edges connect co-expressed genes. Gene names are black for *cell cycle* and red for *plastid fission*. Nodes represent genes, and edges connect co-expressed genes. White nodes indicate genes that were not significantly changed or not detected; gradient-colored nodes represent DEGs, with color intensity reflecting log₂FC (*IE KEA* mutant vs WT). ML DEGs were used for Chlamydomonas. **(B)** Primary and secondary co-expression interactions between *plastid fission* and *cell cycle* genes found in the networks. Co-regulated downregulation is based on both genes being downregulated DEGs. Cut-off |log_2_FC| ≥ 0.75. **(C)** Comparative co-expression network analysis of a *cell cycle* clusters. Nodes represent genes, solid edges connect co-expressed genes and dashed edges connect orthologues. Coloured shapes represent different orthogroups. Cluster comparisons were based on Jaccard Index (JI) computation. See Supplemental Table S6 and S10 for Gene IDs and descriptions.

Interestingly, in Arabidopsis co-expression networks, *cell cycle* genes show little to no regulatory concordance with *plastid fission* genes. Transcripts of *ARC5*, *PDV1*, and *PDV2* displayed log_2_FC values between -0.75 and 0, while *FTSZ2-1*, *FTSZ2-2*, *ARC6*, *MSL2*, *MSL3*, *MCD1*, *KASI*, *MURE*, and *AT2G21280* (*GC1*) ranged between 0 and 0.75, indicating that regulating these genes in response to the loss of IE *AtKEA* had little biological significance for Arabidopsis. While certain genes involved in Arabidopsis chloroplast division were regulated with absolute values higher than a |log_2_FC| of 0.75 (ie. upregulated *YLMG1-1* and *FTSHi1*), they were not concertedly downregulated with *cell cycle* genes as in Chlamydomonas.

To further substantiate co-expression between *cell cycle* and *plastid fission* genes, we plotted the percentage of co-expressed interactions (direct and secondary) for each species (Figure 7B). Within the Arabidopsis co-expression network, 5.3% (4 genes) are direct interactors, while 94.7% (72 genes) are secondary interactors. However, none of these genes were strongly (|log_2_FC| ≥ 0.75) differentially expressed in *At-kea1kea2* mutants. For Chlamydomonas, 100% (4 genes) of the genes are involved in secondary interactions, with 75% (*ARC5*, *ARC6, MIND*) showing co-regulated downregulation.

To identify orthologs of *cell cycle* genes in Chlamydomonas and Arabidopsis, we used an orthogonal approach building on CoNekT (Figure 7C). When comparing co-expression clusters of TOPII, we found several *cell cycle* genes were orthologs between species. *ROW1*, *MCM10*, *PAKRP1L*, *POK1*, *POK2*, *ZCF125* were found to have orthologs within Chlamydomonas that were not annotated by best BLAST hits. Within this gene family, three Chlamydomonas genes were grouped together with seven genes. For *plastid fission* genes, no apparent similarity between Chlamydomonas and Arabidopsis clusters were found. This analysis also indicates that the biological process *cell cycle* in Chlamydomonas showed certain similarities with Arabidopsis.

### Single-Cell Time-Lapse Imaging Reveals Cell Cycle irregularities in Cr-kea1

*Cr-kea1* mutant cells displayed pronounced size distribution differences in micrographs and when analyzed by flow cytometry (Figure 3F). In addition, coordinated downregulation of cell cycle and chloroplast division genes was observed in the *Cr-kea1* transcriptome (Figure 4D, E). In Chlamydomonas, cell division proceeds via a multiple fission cycle, in which cells undergo n divisions (S/M phase) into 2^n^ daughter cells after a prolonged G1 phase. The size of the mother cell determines the number of divisions resulting in 2, 4, 8, 16, etc. equally sized daughter cells (Figure 8A; Cross and Umen, 2015).

**Figure 8:**
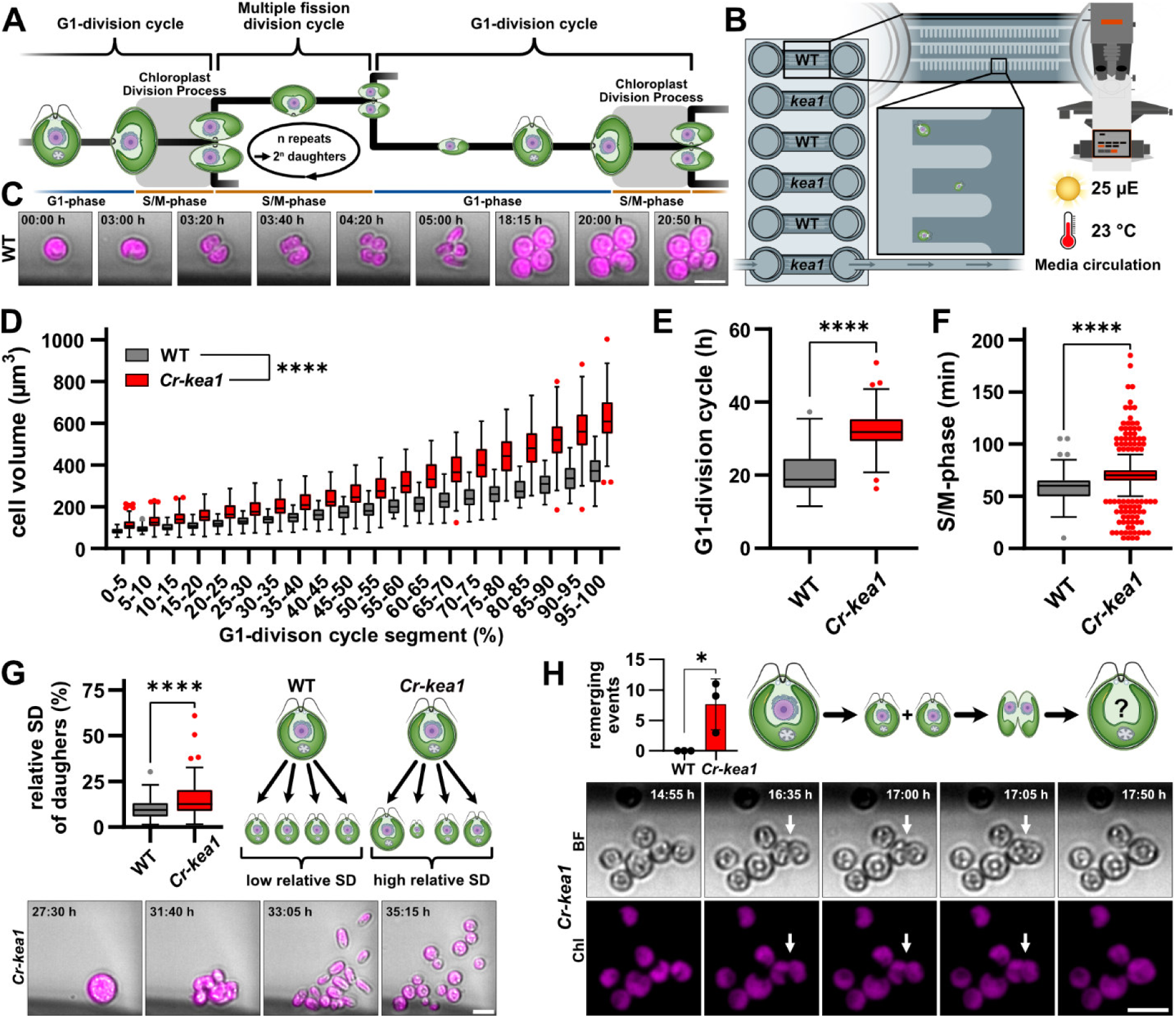
Single Cell Live Imaging reveals cell cycle defects of *Cr-kea1*. **(A)** Simplified diagram of the Chlamydomonas cell cycle, showing the labeling of cell-cycle phases used in this study. **(B)** Schematic of the microfluidic flow-cell chip containing hydrogel-based channels with imprinted cavities that immobilize multiple individual cells for long-term observation. 19 WT- and 18 *Cr-kea1*-mother cells including their first generation of G1-daughter cells were analysed. **(C)** Representative time-lapse image series of WT cells illustrating synchronized growth and division. **(D–F)** Quantitative analysis of single-cell trajectories shows that *Cr-kea1* cells exhibit significantly increased cell volume in every G1-division cycle segment (n^WT^=67; n*^kea1^*=90), prolonged G1 phase (n^WT^=68; n*^kea1^*=90), and extended S/M phase compared to WT (n^WT^=203; n*^kea1^*=633). **(G)** *Cr-kea1* daughter cells display greater size variation following division than WT. Each data point represents the normalized volume difference among daughter cells derived from a single grandmother cell. Daughter cells of n^WT^=55 and n^kea1^=79 grandmothers were compared. **(H)** “Remerging” events, where nearly separated daughter cells fuse again, occur more frequently in *Cr-kea1* and were not observed in WT. The bar graph summarizes data from three replicate channels per genotype. *For all panels:* Representative time-lapse series are shown. Scale bars, 10 µm. Box plots follow Tukey’s convention. Statistical significance was determined with Mann-Whitney tests (D-G) and unpaired t-test (H) and is indicated by asterisks (* p < 0.05; ** p < 0.01; *** p < 0.001; **** p < 0.0001).

The intrinsic heterogeneity of batch cultures poses substantial challenges for the identification of cell cycle stage-specific defects. To overcome these limitations, we developed a live-imaging platform capable of tracking individual cells over several days. Cells were applied to a custom-made, biologically inert hydrogel with multiple chambers embedded in a flow cell chip (Figure 8B). Our design allows for tight temperature- and light-control, constant medium flow, nutrient diffusion, and continuous imaging throughout Chlamydomonas cell division cycles. In addition, the multiple chambers situated on a single chip enabled simultaneous side-by-side growth and monitoring of WT and *Cr-kea1* mutant strains under the exact same controlled culturing conditions in TAP medium at 25 µE (Figure 1G).

WT cells displayed regular and symmetric division with high fidelity, verifying ideal and well-suited growth conditions in our continuous flow cell chip setup (Figure 8C; Supplemental Figure S16A). By contrast, our analysis revealed several key differences in the mutant compared to WT: *Cr-kea1* cells were significantly larger than WT cells during every segment of the G1-division cycle (Figure 8D). Interestingly, the elevated volume of *Cr-kea1* cells can be traced back to the mid-G1 phase (Supplemental Figure S16B), when mutant cells exhibit a significantly higher volume increase than WT. Additionally, mutant cells not only remained longer in the G1 phase (Figure 8E) but also took longer to complete the S/M phase (Figure 8F).

Upon closer examination, specific cytokinesis defects were observed more frequently in *Cr-kea1* mutants: 1) Divisions often produced daughter cells of unequal size (Figure 8G), hinting at a misplaced division plane. Further analysis of mutant daughter cells revealed that these birth volume deviations did not affect subsequent G1-division cycle length differences compared to those of WT daughters (Supplemental Figure S16C). 2) Remerging of seemingly already divided cells multiple hours into the G1 phase was observed in approximately 3% of all *Cr-kea1* divisions but was never detected in WT (Figure 8H). Remerged cells divided into 3 or 5 daughter cells after completing their G1 phase (Supplemental Figure S16A), possibly due to two coexisting nuclei in a single remerged cell.

To further quantify division defects, we determined the fraction of mother cells that failed to produce viable dividing progeny (including rupture, disintegration, or non-dividing daughter cells), which occurred more frequently in *Cr-kea1* (28%, n = 60) than in WT cells (19%, n = 57).

Our single-cell imaging revealed pronounced defects in cell cycle progression and division fidelity, confirming the transcriptomic observations.

## Discussion

Plastid integrity relies on tightly controlled ion fluxes across the inner envelope membrane. Our findings establish CrKEA1 as the functional counterpart of AtKEA1/2 in Arabidopsis, adding experimental support to the hypothesis that KEA-mediated K⁺/H⁺ exchange at the plastid envelope represents an ancient and conserved feature of plastid physiology that likely emerged early during plastid evolution. Loss of *CrKEA1* is associated with alterations in plastid morphology and reduced photosynthetic performance under moderate-light conditions, defects in plastid rRNA maturation, and delayed cell-cycle progression. Together, these phenotypes reveal both conserved and lineage-specific aspects of KEA function across the green lineage (Figure 9).

**Figure 9:**
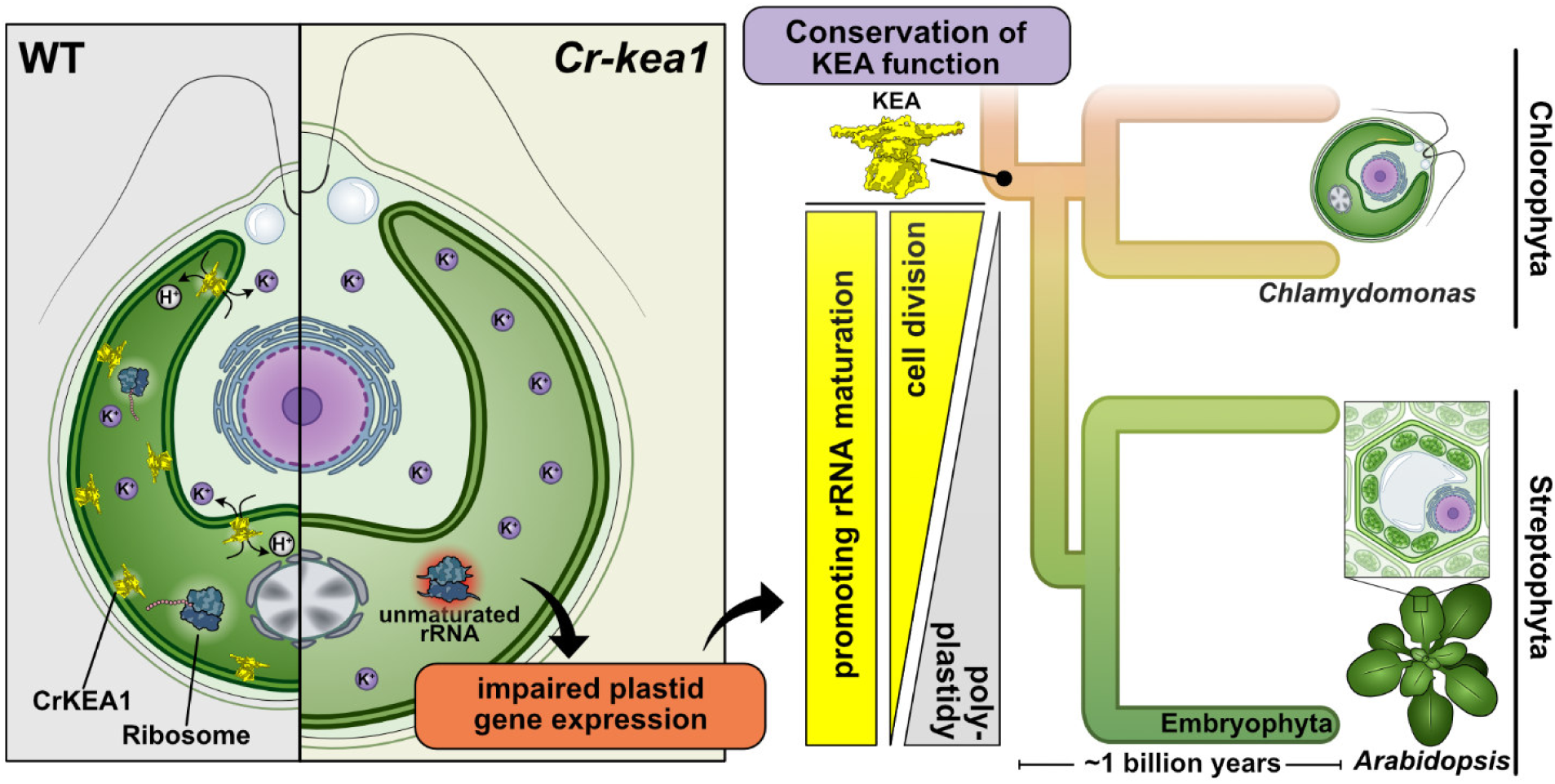
Working Model: Conserved role of inner envelope KEA transporters across the green lineage. In Chlamydomonas, loss of the inner envelope KEA transporters (IE KEAs) leads to K^+^ imbalance and excess stromal K⁺, resulting in swollen, misshapen chloroplasts and impaired rRNA processing required for plastid ribosome biogenesis. IE KEAs, including CrKEA1 in Chlamydomonas and AtKEA1/2 in Arabidopsis, are conserved throughout the green branch of the Archaeplastida, spanning the Chlorophyta (green algae) and Streptophyta (charophytes and land plants). All lineages evolved from a monoplastidic unicellular ancestor characterized by tightly coupled plastid and cell division (*monoplastidic bottleneck*, de Vries and Gould, 2018), a condition still retained in Chlamydomonas. During evolution, the emergence of multicellularity and polyplastidy, as in Arabidopsis, relaxed this coupling and allowed plastid division to proceed independently of the cell cycle. Our findings suggest that the importance of IE KEA-mediated ion homeostasis for error-free cell division diminished with the transition to polyplastidy, while its role in maintaining proper rRNA maturation and ribosome assembly remains conserved across Chlorophyta and Streptophyta.

In Arabidopsis, disruption of *AtKEA1/2* causes increased cellular K^+^ levels, fragile chloroplasts, impaired photosynthesis, and growth (Kunz et al., 2014; Aranda-Sicilia et al., 2016). We observed related phenotypes in *Cr-kea1* mutants, including altered ion balance, growth delay, changes in chloroplast morphology, and reduced photosynthetic efficiency at increased irradiance. These parallels suggest that KEA activity at the plastid envelope plays a broadly conserved role in maintaining the ion homeostasis required for plastid function. The strong conservation of this role is further supported by cross-species complementation, in which CrKEA1 restored growth, photosynthesis, and rRNA processing in *At-kea1kea2* mutants. However, it remains challenging to distinguish primary consequences of impaired plastid ion transport from downstream physiological responses triggered by altered photosynthetic performance and metabolic status.

Despite this limitation, our findings accentuate the potentially fundamental role of IE KEAs in supporting cellular biochemistry by maintaining pH and ion homeostasis in phototropic eukaryotes — a function that may have been conserved since the last common ancestor, likely persisting through more than a billion years of evolutionary divergence (for molecular clock, see Strassert et al., 2021 and Bierenbroodspot et al., 2025).

Consistent with this deep evolutionary conservation are the parallels between plants and algae in the role of KEA transporters in plastid rRNA maturation. In Arabidopsis, loss of AtKEA1/2 disrupts rRNA processing and plastid translation (DeTar et al., 2021). Similarly, *Cr-kea1* mutants accumulated unprocessed 23S-derived precursors, in accordance with impaired maturation. Although the 23S rRNA maturation pathways differ between algae and plants — Chlamydomonas generates unique 7S and 3S fragments, whereas Arabidopsis produces a separate 4.5S rRNA from the opposite end of the 23S precursor — the accumulation of immature rRNA in *Cr-kea1* closely mirrors the maturation defects seen in *At-kea1kea2* (DeTar et al., 2021; DeTar et al., 2022). Structural work offers a mechanistic basis for this observation, showing that K^+^ and magnesium ions are required to stabilize rRNA and the large ribosomal subunit (Nierhaus, 2014; Rozov et al., 2019; Klein et al., 2004). Ion imbalance in IE *KEA* mutants could potentially destabilize rRNA precursors and interfere with their proper processing, or indirectly affect maturation through feedback from defective ribosome assembly (DeTar et al., 2021). Together, our findings strengthen the model that ion imbalance destabilizes rRNA secondary structures and disrupts ribosome biogenesis across systems, despite lineage-specific differences in processing outcomes.

Transcriptomic profiling revealed a conserved upregulation of ribosome biogenesis genes and downregulation of PhANGs in *Cr-kea1*, mirroring the *At-kea1kea2* response (DeTar et al., 2021). In both species, repression of photosynthetic gene expression likely reflects a protective adjustment to plastid dysfunction. In *At-kea1kea2* mutants, the retrograde response is a GUN1-mediated GOLDEN2-LIKE (GLK1/GLK2) transcription factor suppression to delay chloroplast biogenesis, thereby minimizing photo-oxidative damage until stress has been resolved (DeTar et al., 2021). However, this regulatory logic is unlikely to apply to Chlamydomonas: 1) GLK1/2 emerged as a hub controlling the genetic networks that underpin chloroplast differentiation and photomorphogenesis only in zygnematophytes (Dadras et al., 2023), and 2) although GUN1 is evolutionarily ancient, its role in retrograde signaling is thought to have arisen after streptophyte algae diverged from chlorophytes (Honkanen and Small, 2022). Thus, PhANG suppression in Chlamydomonas must rely on different upstream cues. In *Cr-kea1*, where we also found induced ROS-responsive transcripts, an upregulation of *PCYA*, a bilin-synthesis gene, points to tetrapyrrole-derived retrograde signals to repress PhANGs during plastid stress as a coping mechanism (Duanmu et al., 2013).

Because the compared datasets were generated under different experimental contexts and in distinct organisms, these similarities should be interpreted as indicative of broadly conserved stress responses rather than direct regulatory equivalence.

The concurrent induction of Na⁺/SO₄²⁻ (SLT1–3;Pootakham et al.,2010) and K⁺ transporters (particularly *HAK*, *NAK;* Li et al., 2023) together with the altered expression of other ion pumps, exchangers and channels, suggests that disruption of plastid K⁺/H⁺ exchange triggers broader adjustments in cellular ion homeostasis. These regulatory patterns may reflect systemic compensatory responses to altered ion gradients or membrane potentials resulting from impaired K⁺/H⁺ exchange at the plastid envelope. Increased sulfate uptake could further support glutathione biosynthesis, mitigating oxidative stress associated with disturbed ion balance (Niyogi, 1999; Zhang et al., 2004). Also, the tonoplast K⁺ channel *KCN11* was slightly upregulated. Interestingly, *kcn11* mutants display growth defects under nitrogen limitation, including enlarged cell size (Xu and Pan, 2020; Fauser et al., 2022; Vilarrasa-Blasi et al., 2024). These findings point towards the functional coupling of K^+^ and sulfate fluxes as part of the wider physiological consequences of disrupted IE KEA-mediated ion transport in Chlamydomonas. In addition to these conserved features, IE *CrKEA* loss in Chlamydomonas exhibited lineage-specific consequences. *Cr-kea1* mutants showed constitutive induction of the MARS1-dependent chloroplast unfolded protein response (cpUPR; Perlaza et al., 2019). Unlike Arabidopsis, where cpUPR markers were absent or even downregulated (e.g., *CLPB3*) in *At-kea1kea2* (DeTar et al., 2021), Chlamydomonas activates these stress pathways constitutively (already under low light), consistent with destabilized stromal protein homeostasis as soon as ion balance is disrupted by loss of *CrKEA1*. This points to a lineage-specific strategy: whereas multicellular plants can attenuate plastid protein synthesis and postpone chloroplast biogenesis to avoid protein aggregation, the single-celled alga Chlamydomonas cannot completely postpone chloroplast function without jeopardizing survival. Thus, the unicellular alga must maintain translation in its single essential plastid and instead relies on cpUPR pathways to counteract misfolding and aggregation (Dogra et al., 2019; Perlaza et al., 2019; Richter et al., 2023).

Further characteristic observed in *Cr-kea1* mutants were enlarged cells, an extended G1 phase, irregular cytokinesis, and strong downregulation of cell division genes. Both *CDKB1* and its partnering cyclins (*CYCA1*, *CYCB1*) were strongly downregulated, consistent with defects in mitotic progression and multiple fission cycles (Tulin and Cross, 2014; Bisova et al., 2005). The partial reduction of D-type cyclins further weakens G1/S entry, while the mild repression of CDKG1 implicates misregulation of size-dependent checkpoints (Li et al., 2016; Umen and Goodenough, 2001). The coupled downregulation of plastid fission genes that we found through co-expression network analyses aligns with the tight synchronization between plastid division and cell cycle progression in unicellular algae (Sumiya et al., 2016; Clark-Cotton et al., 2025). It was shown that conditional repression of plastid genes (*rpoA*, *rps12*) that are required for the expression machinery halts Chlamydomonas growth whereas cells resume proliferation once chloroplast function is restored (Ramundo et al., 2013). The cell cycle in *Cr-kea1* is not arrested. Instead, mutant cells progress more slowly through G1, delaying the onset of cell and chloroplast division. After entry into S/M phase, cells more often exhibit specific division defects, including potentially misplaced division planes and incomplete cytokinesis, which may lead to remerging events. Elevated light intensity, which stimulates growth and cell division in algae (Dupuis et al., 2025), was associated with an increased occurrence of enlarged cells, consistent with division defects. However, we did not assess whether the frequency of defects changes under these conditions.

In contrast to the cell cycle-coupled plastid division in Chlamydomonas, plant plastids divide autonomously, controlled by developmental programs, tissue context, and land-plant-specific proteins such as PDV1/2 (Okazaki et al., 2009; Vercruyssen et al., 2015). Thus, while IE *KEA* loss in both systems impairs plastid function, the downstream impact is filtered through fundamentally different organizational logics: cell cycle-plastid coupling in algae versus tissue-specific developmental regulation in plants. However, our finding that *Cr-kea1* cells downregulate plastid division and accumulate enlarged chloroplasts parallels observations in Arabidopsis, where *At-kea1kea2* mutants harbor fewer but enlarged plastids per cell (Kunz et al., 2014; Aranda-Sicilia et al., 2016). Despite differences in cellular architecture — one plastid per algal cell versus many plastids per plant cell — KEA transporters thus appear broadly required to coordinate plastid division and maintain organelle size.

At present, it remains difficult to distinguish whether the observed cell-cycle defects arise directly from perturbed ion homeostasis or represent secondary consequences of reduced photosynthetic performance and metabolic stress. In Chlamydomonas, cell-cycle progression is tightly coupled to cellular energy status. Moreover, photoinhibition or chloroplast dysfunction is likely to delay progression through G1. Thus, the enlarged cell size and delayed division observed in *Cr-kea1* may reflect a combination of direct plastid signaling effects and indirect responses to altered photosynthetic capacity.

While the present work defines key physiological consequences of *CrKEA1* loss, future work will be required to disentangle primary ion transport defects from downstream physiological consequences. In particular, compartment-resolved measurements of K^+^ and pH, for example using targeted fluorescent sensors or electrophysiological approaches, would help clarify how CrKEA1 activity shapes ion homeostasis. Combining such approaches with time-resolved ionomics across the diurnal cycle, and their extension to single-cell measurements, may further reveal how perturbations in chloroplast ion transport propagate to whole-cell physiological responses and cell-cycle dynamics.

## Conclusions

Given that K^+^ fluxes have universal roles in driving cell-cycle progression in microalgae and beyond (Urrego et al., 2014; Feeney et al., 2016; Henslee et al., 2017; Rodríguez et al., 2024), our integrated ionomics-single-cell imaging approach offers a roadmap for future studies to unravel how perturbations in organellar ion transport translate into systemic defects in growth, proliferation, and stress adaptation. Taken together, our results support a model in which IE KEA transporters are evolutionarily conserved regulators of plastid homeostasis, linking ion transport to ribosome assembly and photosynthetic capacity. Yet the downstream phenotypes diverge, reflecting the distinct cellular architectures of unicellular algae and multicellular plants. *Cr-kea1* mutants expose defects in cell cycle fidelity absent in plants, while Arabidopsis mutants reveal developmental and signaling adjustments not observed in algae. These findings illuminate how plastid ion transport serves as a deeply conserved module, whose consequences for cell physiology depend on lineage-specific evolutionary trajectories.

## Material and Methods

### Algae cultivation and growth measurements

All Chlamydomonas (*Chlamydomonas reinhardtii*) *kea1* mutants and complementation lines generated in this study share the CC-406 cw15 mt- background (Davies and Lyall, 1973) which served as the reference (“wild type”, WT) in this study.

Algae were grown in Tris-acetate-phosphate (TAP; Gorman and Levine, 1965) medium under constant light of 25 µmol photons m^-2^ s^-1^ at 23°C and shaking at 125 rpm if in liquid culture, unless indicated otherwise. TAP was supplemented with 1% (w/v) sorbitol for culturing strain cc406. For photoautotrophic conditions algae were grown in Sueoka’s high salt medium (HSM; Sueoka, 1960).

The optical density at 750 nm (OD750) of liquid cultures was determined by a UV-vis spectrophotometer and the cell count ml^-1^ by a BD Accuri^TM^ C6 plus flow cytometer. To determine a conversion factor to calculate the size of the cells measured from FSC-A, the flow cytometer was calibrated with CS&T RUO Beads (BD Biosciences, USA). The derived linear regression was y = 2.432 x10^6^ x + 1,706 with y being the diameter (µm) and x the FSC-A values.

### Plant growth and generation of CrKEA1-YFP plants

Arabidopsis (*Arabidopsis thaliana*) plants were grown at 110 µE illumination in long-day conditions (16 h/8 h at 22°C). To generate transgenic *CrKEA1* Arabidopsis plants, the codon-optimized *CrKEA1* coding sequence (Cre04.g220200_4532.1, fused to a Twin-Strep tag) was cloned into the binary Ti vector *pG20_TEV_mVenus_Hyg* via Golden Gate assembly downstream of the UBQ10 promoter (Addgene ID #240651; Schwartz et al., 2025) as described in Supplemental Table S11. Venus is referred to throughout as YFP. The resulting *CrKEA1-TwinStrep-TEV-YFP* (short *CrKEA1-YFP*) fusion construct was used for stable transformation of the Arabidopsis T-DNA insertion line *At-kea1*-*1kea2*-*1* (Kunz et al., 2014). The final construct was transformed into Arabidopsis via the *Agrobacterium tumefaciens* (GV3101 containing *pSOUP*) by the floral dip method as described previously (Clough and Bent, 1998). The T_1_ generation was selected on 25 µg/ml Hygromycin-containing 1/2 strength MS medium (Murashige and Skoog, 1962) and screened for phenotypical rescue of the *At-kea1*-*1kea2*-*1* mutant. Expression of *CrKEA1-YFP* was verified via immunoblotting (Höhner et al., 2019).

### Generation of Chlamydomonas kea1 mutants by CRISPR/Cas9

*Cr-kea1* mutants were created in CC-406 cw15 mt⁻ using insertional mutagenesis by the CRISPR/Cas9 technique following the protocol established (Greiner et al., 2017). Briefly, a crRNA sequence was designed to target *CrKEA1* exon-19. The crRNA was annealed with tracrRNA (Integrated DNA Technologies, #1072533) to generate guide RNAs (gRNA) and then conjugated with Alt-R® S.p. Cas9 Nuclease V3 (Integrated DNA Technologies, #1081058) to form the SpCas9/gRNA complex. The latter was co-transformed into Chlamydomonas with a mutation oligonucleotide containing orientation and frameshift tolerant STOP codons and with the plasmid pBC1 (pJR38, Neupert et al., 2009) conferring paromomycin resistance. Transformation was carried out on a NEPA21 electroporator (Nepa Gene). Cells were selected under 10 µg ml^-1^ paromomycin and screened by colony-PCR. Knock-out of *CrKEA1* was confirmed by immunoblotting using CrKEA1 antibodies. Relevant sequences are listed in Supplemental Table S1.

### Generation of Cr-kea1 complementation lines

The *CrKEA1* gDNA sequence, including the native promoter, was inserted into *pLM161* (Cr-Venus-3xFLAG; hygromycin), applying the recombineering approach described in Emrich-Mills et al., 2021; Supplemental Table S12). Nuclear transformation was achieved by the glass bead method (Kindle, 1990). Transformed *Cr-kea1* cells were selected under 100 µg ml^-1^ hygromycin followed by screening for the expression of the CrKEA1-YFP fusion protein by fluorescence microscopy and immunoblot.

### Design of the CrKEA1 antibody

To generate antibodies against CrKEA1, a DNA fragment codon optimized for *E. coli* synthesized by Twist Bioscience (California, USA), was cloned into the plasmid *pHue* (Catanzariti et al., 2004; modified for Gold Gate cloning) resulting in the plasmid *pHue_CrKEA1-aa94-337* (Supplemental Table S13). Proteins were expressed in *E. coli* strain BL21 (DE3) overnight at 23°C and lysed in lysis buffer (20 mM Tris-HCl, pH 8.0, 300 mM NaCl and 10 mM imidazole). The HisUbiquitin-tagged *CrKEA1* fragment was extracted by an IMAC column, the tag was cleaved by Ubiquitin Specific Peptidase 2 (USP2; Baker et al., 2005) and the sample passed over a size exclusion chromatography (SEC) column in SEC buffer (20 mM Tris-HCl, 300 mM NaCl). The SEC elution was reapplied to an IMAC column, where the tag-less protein was recovered from the flow-through. The pure protein CrKEA1-aa94-337 in 20 mM Tris pH 8, 200 mM NaCl, and 5% glycerol served BioGenes (Berlin, Germany) as antigen to produce polyclonal antibodies. Serum was used for immunodetection.

### Protein extraction and immunoblotting

For Chlamydomonas, cultures grown to an OD750 of 2 were pelleted and resuspended in Laemmli SDS loading dye containing 50 mM DTT. Samples set to an OD of 22 were heated to 60°C for 20 min and then subjected to SDS PAGE. Immunoblotting was performed on PVDF membranes with secondary antibodies coupled to horse radish peroxidase and the enhanced chemiluminescence (ECL) system. For Arabidopsis, immunoblotting was performed as described before (Höhner et al., 2019). The following antibodies were used: GFP (Roche 11814460001), NAB1 (Agrisera AS08 333), AtKEA1 (Bölter et al., 2020), CrKEA1 (this study)

### Element determination via TXRF

Cultures for element analysis were washed thrice in isotonic buffer (300 mM sorbitol, 20 mM HEPES pH 7.5, 5 mM EDTA, 10 mM NaHCO_3_). Cell numbers in washed samples were individually determined for each sample using a Neubauer Improved counting chamber (Hecht-Assistent, Sondheim vor der Rhön, Germany). Elemental analysis was performed by total reflection X-ray fluorescence (TXRF) using an S4 T-STAR (Bruker, Berlin, Germany) as described in Holzner et al., 2026. The sample was mixed in a 1:1 ratio with internal standards to final concentrations of 1 ppm vanadium (W-L) and 1 ppm gallium (Mo-K) and 0.1% (w/v) polyvinyl alcohol (PVA). Samples were applied to silicon-coated quartz glass carriers and measured at 50 kV for 1000 s per sample using Mo-K and W-L excitation. To determine background element levels, a sample of isotonic buffer was measured, which was subtracted from the respective samples. Element concentrations were normalized to the cell number of each sample. Where indicated, values were additionally normalized to the average cell volume as determined by flow cytometry (FC).

### Determination of cytosolic pH

Cytosolic pH was determined using the ratiometric fluorescent dye BCECF-AM, based on the approach described previously (Pang et al., 2023). Cells were harvested and resuspended to ∼2 × 10⁷ cells ml⁻¹ in NMG buffer (10 mM HEPES, 60 mM KCl, 3 mM MgCl₂, pH 6.8). BCECF-AM was added to a final concentration of 5 µM, and cells were incubated for 1 h at 36 °C to allow dye uptake and intracellular ester cleavage. Cells were then washed and resuspended in medium to remove extracellular dye. Fluorescence measurements were performed with a Spark plate reader (TECAN, Switzerland) in 96-well plates using excitation wavelengths of 440 ± 10 nm and 490 ± 10 nm and emission detection at 535 ± 10 nm. Dye uptake and localization were verified with confocal imaging (Stellaris 5, Leica, Wetzlar). BCECF fluorescence was excited at 448 nm and emission was detected at 530 ± 10 nm.

For calibration, TAP was substituted with NMG buffer (10 mM HEPES, 60 mM KCl, 3 mM MgCl₂, pH 6.8). BCECF-loaded cells were equilibrated with NMG buffers of defined pH (4–9) containing 100 µM nigericin to equilibrate intra- and extracellular pH. The resulting titration curve (average of three replicates) was used to convert fluorescence ratios (490/440 nm) to cytosolic pH values.

### Pulse-Amplitude-Modulation (PAM) fluorometry and leaf area

IMAGING-PAM systems (M-Series Maxi and Hexagon; Walz GmbH, Effeltrich, Germany) were used for leaf area and PAM measurements. For Arabidopsis, experiments were performed as described previously (Schneider et al., 2019). For Chlamydomonas drop cultures, spotted cultures were cultivated under standard conditions for 3 days before subjecting them to PAM measurements. For liquid culture, cells grown in 50 ml were adjusted to an OD of 1.5, and equal volumes (2 ml) were transferred into multi-well plates for PAM measurements after 15 min dark adaptation. For NPQ induction and photoinhibition (PI) experiments, cultures were adjusted to OD₇₅₀ ≈ 0.12 and transferred to 10 ml beakers that were continuously stirred during measurements. NPQ induction was measured under actinic light (646 PAR) following standard PAM protocols. For photoinhibition experiments, cultures were exposed to actinic high light (646 PAR) and recovery of PSII efficiency was monitored over time. During recovery, samples were incubated for 20 min under low light (15 µmol photons m⁻² s⁻¹), followed by 10 min dark adaptation before determination of *F*_v_/*F*_m_. This cycle was repeated for 5 h. Lincomycin was added to one set of samples after the first post-illumination *F*_v_/*F*_m_ measurement to inhibit chloroplast protein synthesis.

### Protoplast isolation and localization of CrKEA1-YFP in Arabidopsis and Chlamydomonas

Protoplasts were isolated from Arabidopsis leaves as described before (Wu et al., 2009). Localization of CrKEA1–YFP in Arabidopsis protoplasts and intact Chlamydomonas cells was performed using a Stellaris 5 confocal laser scanning microscope (Leica, Wetzlar, Germany), equipped with a 405 nm diode laser, a supercontinuum white light laser (WLL), and a Power Hybrid (HyD S) detector. Chlorophyll autofluorescence was excited at 405 nm and detected between 620 and 810 nm, while YFP was excited at 515 nm and detected between 520 and 580 nm.

### Transmission Electron Microscopy

Sample preparation was performed as previously described (Lübben et al., 2024). Chlamydomonas samples were frozen under high pressure (2,100 bar) in culture medium using an EM HPM100 (Leica Microsystems, Germany). Following cryofixation, samples underwent freeze substitution (Peschke et al., 2013; Walther and Ziegler, 2002) in an EM AFS2 system (Leica Microsystems), after which they were embedded in Epon 812 and polymerized for 16 h at 63 °C. Ultrathin sections of 50 nm (ultra 35°, 3.0 mm, DiATOME) were cut on an Ultracut E ultramicrotome (Leica Microsystems) and collected on collodion-coated, 400-mesh copper grids (Science Services GmbH, Germany). Prior to imaging, sections were post-stained with lead citrate (Reynolds, 1963). Samples were examined in a Zeiss EM 912 transmission electron microscope (Zeiss, Germany) equipped with an integrated OMEGA energy filter operated in zero-loss mode at 80 kV. Images were captured at a nominal magnification of 12,500× using a 2k×2k slow-scan CCD camera (Tröndle Restlichtverstärkersystem, Germany).

### Whole Genome Sequencing

gDNA was isolated according to Dutcher et al., 2012 and sent to BMKGene (Biomarker Technologies, Münster, Germany / Hongkong) for Illumina sequencing. CLC Genomics Workbench and Integrative Genomics Viewer (IGV) was used to confirm stop codon insertion in exon 19. TDNAscan (Sun et al., 2019) was used to search for additional off-target insertions.

### RNA-seq after ribo-depletion, Gene Ontology analysis, and constructing co-expression networks

Four replicates per genotype and condition were sampled to obtain RNA-seq from the light shift experiment. RNA was extracted using TRIzol™ reagent (Ambion, USA; Simms et al., 1993) following the manufacturer’s instructions. RNA was further purified and DNA contaminations were removed using the Monarch® RNA Cleanup Kit (NEB) according to the manufacturer’s protocol. RNA quality control, depletion of ribosomal RNA, library preparation, and eukaryotic lncRNA Sequencing was performed by BMKGene on an Illumina platform, generating over 10 Mio paired-end reads (150 bp) per sample. RNA-seq data were analyzed on the Galaxy platform (usegalaxy.eu; Galaxy-Community, 2024) using the tools, reference genome and annotation file (JGI Phytozome) listed in Supplemental Table S14. For statistical analysis and assessment of differentially expressed genes (DEGs), the low read count filters were set to “1-count-per-million” (CPM=1) in at least 3 samples. DEGs were considered significant with an adjusted p-value < 0.05. Gene Ontology (GO) term enrichment was assessed with g:Profiler (Kolberg et al., 2023) and condensed with REVIGO (Supek et al., 2011). Co-expression networks were calculated using the StreptoNet (https://rshiny.gwdg.de/apps/streptonet/) framework (Dadras et al., 2023) and constructed based on gene lists of interest. Chlamydomonas genes were annotated using the best BLAST hits against Arabidopsis genes. CoNekT (https://conekt.sbs.ntu.edu.sg/) framework was used to compare clusters of co-expression networks. See Supplemental Table S14 for further details.

### RNAseq with focus on ribosomal RNA (MiSeq NGS)

To analyze plastid rRNA processing, RNA was extracted from cells grown in HSM under moderate light (100 µmol m^-2^ s^-1^) as mentioned above. Library preparation was performed using the NEBNext® Ultra™ II RNA Library Prep Kit (NEB). RNA and library quality were assessed using the Agilent 2100 Bioanalyzer. Next-generation sequencing (2 × 300 paired-end) was carried out on a MiSeq sequencer (Illumina, San Diego, CA, USA), generating over 1.5 million paired-end reads per sample with a length of 200-250 nt. Data processing was performed on usegalaxy.eu (Galaxy-Community, 2024) as described above. The effective genome size (113,516,734 bp) was determined by subtracting the total number of ambiguous (N) bases (1,114,981 bp) from the actual Chlamydomonas genome size (114,631,715 bp). To analyze coverage differences between samples, mapped reads were processed through the BamCoverage and BamCompare pipelines of the deepTools suite (Ramírez et al., 2016), focusing on plastome rRNA transcripts. BigWig output files generated from both tools were exported as bedGraph (.csv format).

### Non-Radioactive RNA blots

RNA blots were performed using 3’-Digoxigenin (DIG)-labeled DNA probes from metabion international AG, Germany (Supplemental Table S15; Holtke & Kessler 1990; Ramkissoon et al., 2006). RNA samples were denatured at 55°C for 45 minutes in denaturing buffer (15% (v/v) DMSO, 7% (w/v) Glyoxal in 20 mM MOPS, 5 mM sodium acetate, 1 mM EDTA, pH 7) before adding loading dye. Samples were separated on a 1.2% agarose MOPS gel at 4°C, and transferred onto a nylon membrane via capillary blotting in 20x SSC (3M NaCl, 0.3M Trisodium Citrate dihydrate, pH 7) overnight, followed by UV crosslinking. To develop the blot, the DIG Northern Starter Kit (Roche, Cat. No. 12 039 672 910) was utilized as described by the manufacturer but using 50°C for hybridisation.

### Single-cell time-lapse imaging and image analysis

The microchannel layout was based on the original *Mother Machine* design for bacterial single-cell studies (Wang et al., 2010) and was modified to enable the observation of Chlamydomonas under hydrogel-mediated confinement. Hydrogel patterning was carried out using the photolithographic microfabrication workflow previously described for PEG-based hydrogels (Stöberl et al., 2023), with adjustments to the precursor composition and pattern geometry. Flow channels were filled with diluted cultures (5 × 10^4^ cells mL⁻¹) and placed in a stage-top chamber (okolab, H101-CRYO-BL) connected to a cryostatic water bath (Lauda, RE415S). The medium temperature was adjusted to 23°C and flow rate to 200 µl h^-1^. Cells were continuously illuminated with 25 µE. Images were acquired every 5 min on a Leica DMi8 Thunder microscope (Leica, Wetzlar) using LASX. Chlorophyll autofluorescence was excited with a Leica LED8 at 475 nm and detected with a Leica DFC9000 GTC camera with a 642/80 filter. Image analysis was conducted in Cell-ACDC (Padovani et al., 2022). A custom-trained Cellpose (Pachitariu and Stringer, 2022) model was used to segment cells. Segmentation was controlled and corrected in the Cell-ACDC GUI. Segmented objects were then tracked with Trackastra (Gallusser and Weigert, 2025) with the general_2d model using the chlorophyll autofluorescence channel as input. Truthful tracking was checked in the Cell-ACDC GUI. Out-of-focus cells, division failures and cells with an unobservable future were manually marked in the Cell-ACDC GUI for filtering in downstream analysis. Resulting .csv files were filtered and analysed in a custom phyton script.

### Sequence alignment, structure prediction, and domain annotations

Multiple Sequence alignment was performed by the *MultAlin* online tool (Corpet 1988). Structural models were predicted using AlphaFold3 (DeepMind; Abramson et al., 2024). The default parameters were used, and no experimental structural constraints were applied. For model visualization, we used UCSF ChimeraX (Pettersen et al., 2021). For domain annotations, information obtained from Uniprot.org (Q9ZTZ7) and AlphaFold3 prediction was combined.

### Accession numbers and annotations

Chlamydomonas strains CC-406 cw15 mt⁻, CC-4533 cw15 mt⁻, the CLiP line (Li et al., 2016) LMJ.RY0402.187220, and the BAC (well number 19-B-5) encoding the *CrKEA1* (Cre04.g220200_4532.1) gDNA were obtained from the Chlamydomonas Resource Center (University of Minnesota, USA). The strains *Cr-kea1* and *CrKEA1-YFP* (in *Cr-kea1*), generated in this study, are available as CC-6360 *Cr-*kea1-1 mt^-^ and CC-6361 *Cr-*kea1-1_CrKEA1-YFP mt^-^, respectively. The Arabidopsis T-DNA insertion mutant *At-kea1-1kea2-1* (SAIL_586_D02, SALK_045324) (Kunz et al., 2014) is available in the stock centers: ABRC: CS72318 or NASC: N72318. All other Gene IDs and annotations are listed in the Supplemental Table S4, S6 and S10. RNA-seq data generated in this study have been deposited in the NCBI Sequence Read Archive (SRA) under BioProject accession number PRJNA1343327 (Wunder et al., 2025).

### Disclosure on the use of AI-assisted tools

The original draft of this manuscript was written *de novo* by the authors. In addition, ChatGPT was used for language and copyediting assistance. All generated content was carefully reviewed and edited by the authors, who take full responsibility for the final content of the manuscript.

## Supplemental Data

Supplemental Figure S1: Alignment of inner envelope (IE) KEA protein sequences from Chlamydomonas and Arabidopsis.

Supplemental Figure S2: Generation of the CrKEA1 antibody and testing the CLiP line.

Supplemental Figure S3: Genomic characterization of the *Cr-kea1* CRISPR/Cas9 knock-out site and immunoblotting.

Supplemental Figure S4: Genomic characterization of the *Cr-kea1* CRISPR/Cas9 knock-out site (part2).

Supplemental Figure S5: Characterization of independent *CrKEA1–YFP* (*Cr-kea1*) complementation lines.

Supplemental Figure S6: CrKEA1-YFP localizes to the plastid envelope.

Supplemental Figure S7: Confocal microscopy of chloroplast morphology and ultrastructural analysis of *Cr-kea1* cells.

Supplemental Figure S8: Ionomic profiles expressed per cell and calibration of cytosolic pH measurements in *Cr-kea1* and control strains.

Supplemental Figure S9: Scatter plots corresponding to the flow cytometry experiment shown in Figure 3.

Supplemental Figure S10: Additional results related to the RNA-seq experiment.

Supplemental Figure S11: Comparison of plastid rRNA processing pathways in Chlamydomonas and Arabidopsis.

Supplemental Figure S12: rRNA-seq analysis of plastid rRNA processing including the *CrKEA1-YFP* complementation line.

Supplemental Figure S13: RNA blot analysis of plastid rRNA processing in the *Cr-kea1* mutant.

Supplemental Figure S14: Structural comparison of AtKEA1 and CrKEA1.

Supplemental Figure S15: Characterization of the Arabidopsis *kea1kea2* line complemented with *CrKEA1-YFP*.

Supplemental Figure S16: Single cell live imaging of *Cr-kea1* and WT.

Supplemental Tables

Supplemental References

## Acknowledgements

At LMU, we thank Berina Isakovic and Yuling Ma for help in isolating *Cr-kea1* mutant and complementation lines, Andreas Brachmann for supervising the rRNA seq experiment, and Nikolay Kayo Manavski and Jörg Meurer for teaching RNA work, Felix Thoma for support with the initial fluorescence microscopy experiments, Stefan Stöberl and Bettina Bölter for discussions about hydrogel flow cell chips and result interpretations, respectively. We thank Isabell Andree (Heinrich-Heine-Universität Düsseldorf) for helpful advice on PAM measurements. Lastly, we thank Alexander Schober from NTU Singapore for providing the Goldengate compatible pHue vector.

## Author Contributions

T.W. and H.-H.K. conceived research idea and wrote the manuscript. T.W. cloned constructs, isolated algal and plant mutants, performed most experiments, analyzed data, and designed the figures. L.E. performed cytosolic pH experiments, single-cell imaging, analyzed image data, designed figures, and wrote sections. L.J.H. conducted TXRF and analyzed elemental data. C.S. generated plant mutants and helped with element analysis. L.K. and B.S. performed and assisted in RNA and growth experiments. J.F. carried out RNA blot, NPQ, and recovery-after-photoinhibition experiments.. M.A. isolated algal mutants. F.P. and K.M.S. supported analysis of single-cell imaging data. M.O. performed transmission electron microscopy. B.B. assisted with RNA-seq data analysis and visualization. P.G. and L.C.M.M. recombineered plasmids. Z.M.A. and J.dV. performed cross-species gene network analyses. C.L. and J.O.R. designed and fabricated hydrogel flow-cell chips. S.M. provided illustration. J.T.T. and J.N. designed and carried out initial CRISPR/Cas9. All authors assisted in editing the manuscript.

