## Supplemental Figures for "Conserved and Lineage-Specific Roles of KEA-Mediated Ion Homeostasis in *Chlamydomonas*"

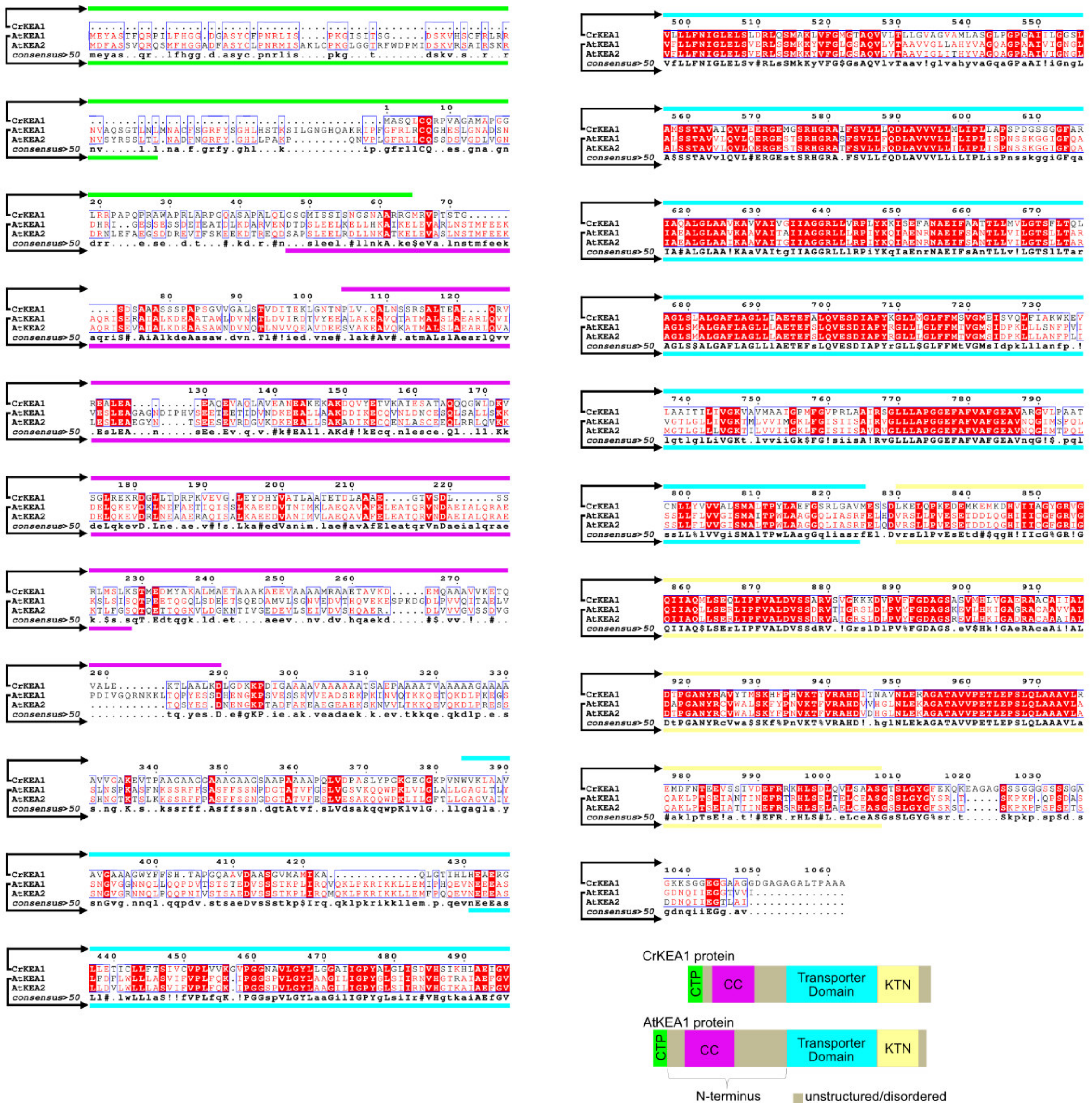


**Figure S1: Alignment of inner envelope (IE) KEA protein sequences from Chlamydomonas and Arabidopsis.** Alignment of IE KEA protein sequences was generated with the MultAlin tool using the following reference sequences: CrKEA1 (Cre04.g220200_4532, Phytozome), AtKEA1 (At1g01790, UniProt Q9ZTZ7), and AtKEA2 (At4g00630, UniProt O65272). Colored highlights below and above the alignment correspond to the simplified domain structure model shown at the lower right, which was guided by AlphaFold3 structure prediction.

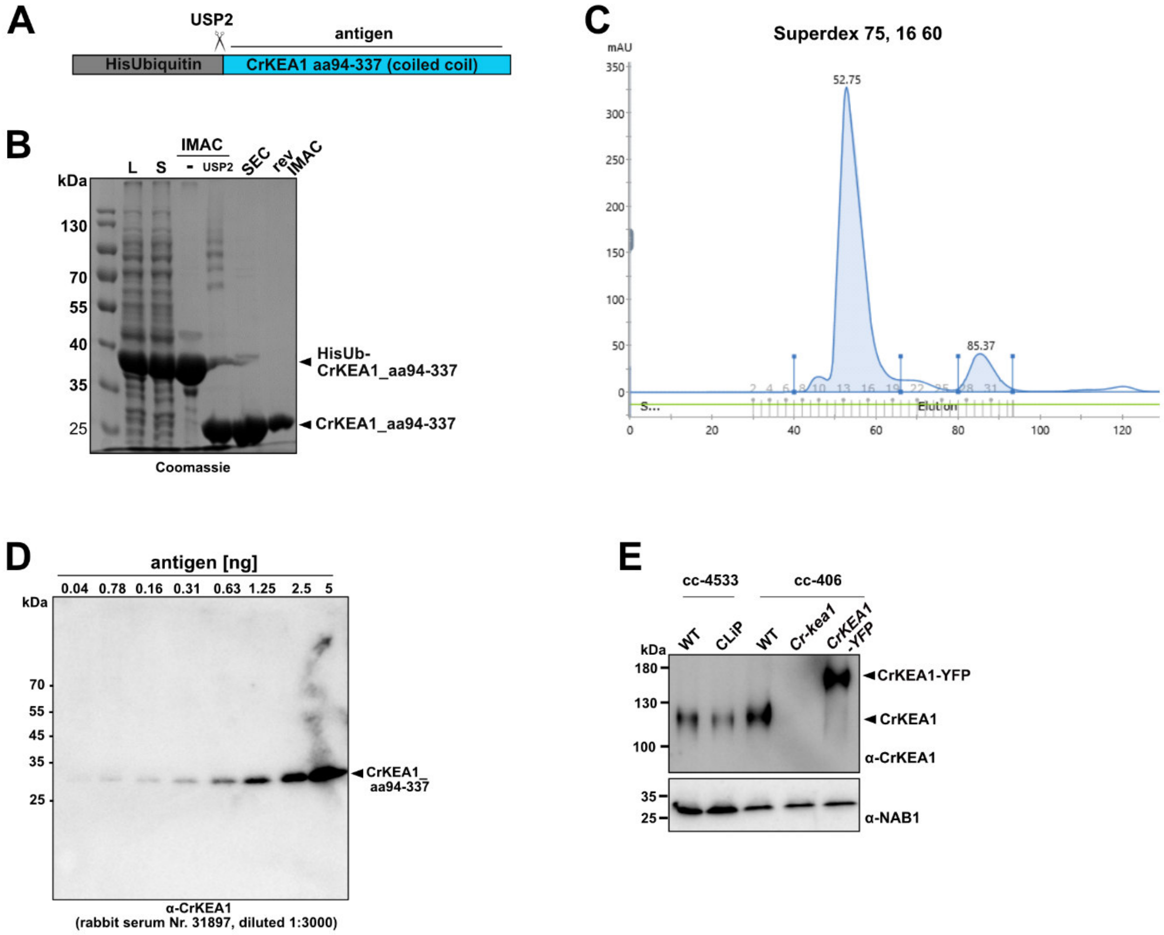


**Figure S2: Generation of the CrKEA1 antibody and testing the CLiP line. (A)** Simplified schematic of the *His*-ubiquitin–tagged CrKEA1 fragment (amino acids 94–337) expressed in *E. coli* for production of the antigen used to generate the CrKEA1-specific antibody. **(B)** Coomassie-stained SDS–PAGE gel showing purification of the antigen from *E. coli* using immobilized metal affinity chromatography (IMAC) followed by size-exclusion chromatography (SEC). The "rev IMAC" fraction was used as the immunogen. Abbreviations: L, total lysate; S, soluble lysate; IMAC, eluted fraction after immobilized metal affinity chromatography; IMAC USP2, IMAC fraction following cleavage of the $His_6$–Ub moiety; SEC, combined peak fractions after size-exclusion chromatography; rev IMAC, flow-through fraction from reapplication to IMAC. **(C)** SEC chromatogram corresponding to the purification shown in panel B. **(D)** Immunoblot using the newly generated *CrKEA1* antibody and serial dilutions of purified antigen. **(E)** Immunoblot of total protein extracts from the potential *CrKEA1* (Cre04.g220200) CLiP knock-out line LMJ.RY0402.187220 compared with the *CrKEA1* knock-out line (*Cr-kea1*) generated in this study via CRISPR/Cas9. The CLiP library line is based on the *cc-4533* "wild-type" background, whereas *cc-406* was used in this study.

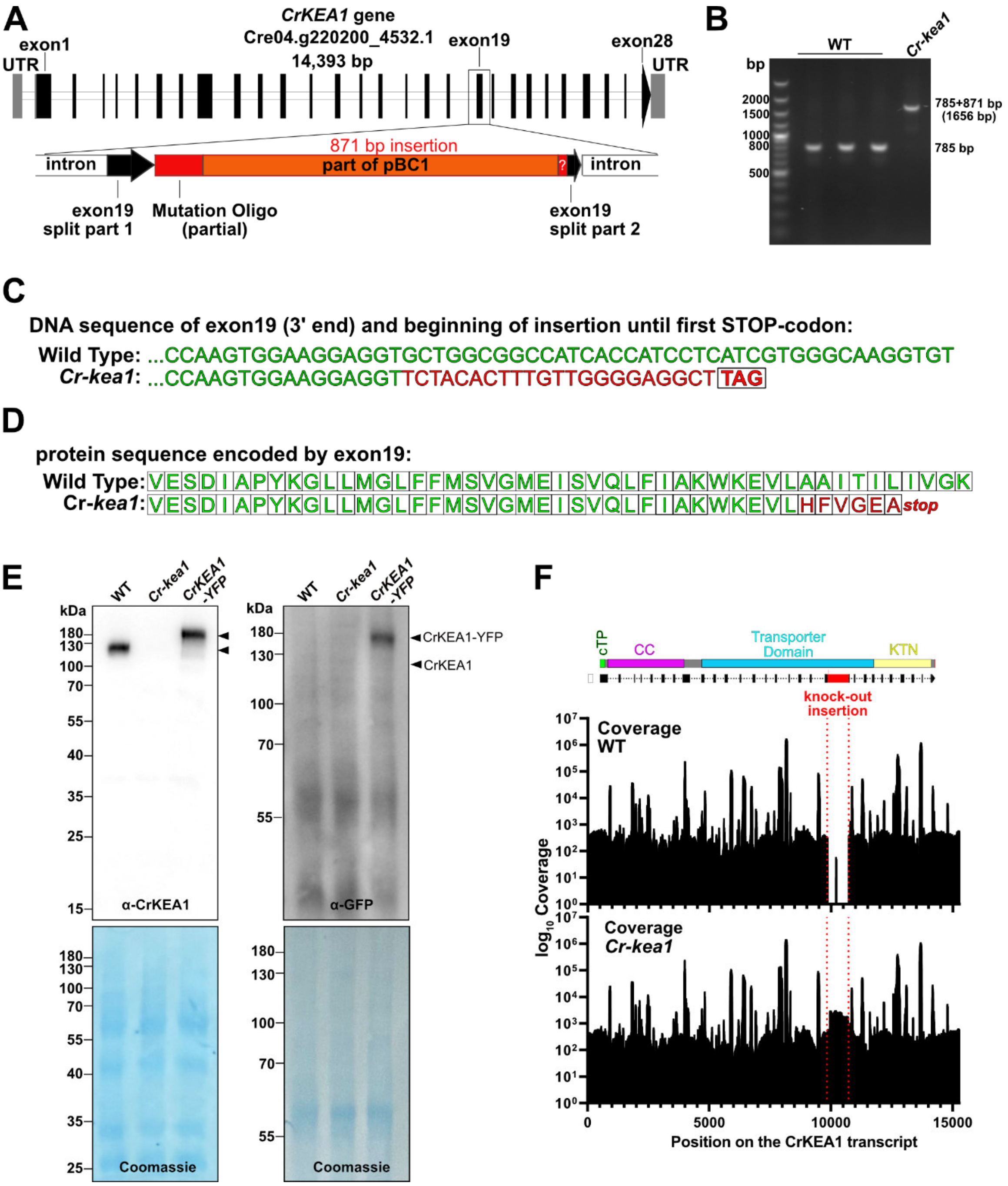


**Figure S3: Genomic characterization of the *Cr-kea1* CRISPR/Cas9 knock-out site and immunoblotting. (A)** Schematic representation of the *CrKEA1* gene showing intron–exon organization, domain structure, and the position of the CRISPR/Cas9-induced insertion which disrupts exon 19. The insertion consists of the mutation oligonucleotide (partial) containing an in-frame stop codon together with a large fragment of the co-transformed plasmid pBC1. **(B)** PCR on genomic DNA extracted from wild type and *Cr-kea1*. Primer binding sites are located in gene-specific regions flanking the disrupted locus. **(C)** DNA sequence comparison between wild type and *Cr-kea1* at the 3′ end of exon 19, including the junction with the inserted fragment up to the first introduced stop codon. **(D)** Comparison of the predicted protein sequence encoded by exon 19 in wild type and *Cr-kea1*, illustrating the premature truncation of the CrKEA1 protein. **(E)** Full immunoblot corresponding to the cropped version shown in Figure 1, including the matching anti-GFP western. **(F)** NGS read coverage across the *CrKEA1* gene aligned to the *CrKEA1* gene model, confirming disruption of exon 19 and loss of intact sequence continuity. cTP, chloroplast targeting peptide; CC, coiled coil domain; KTN, regulatory domain.

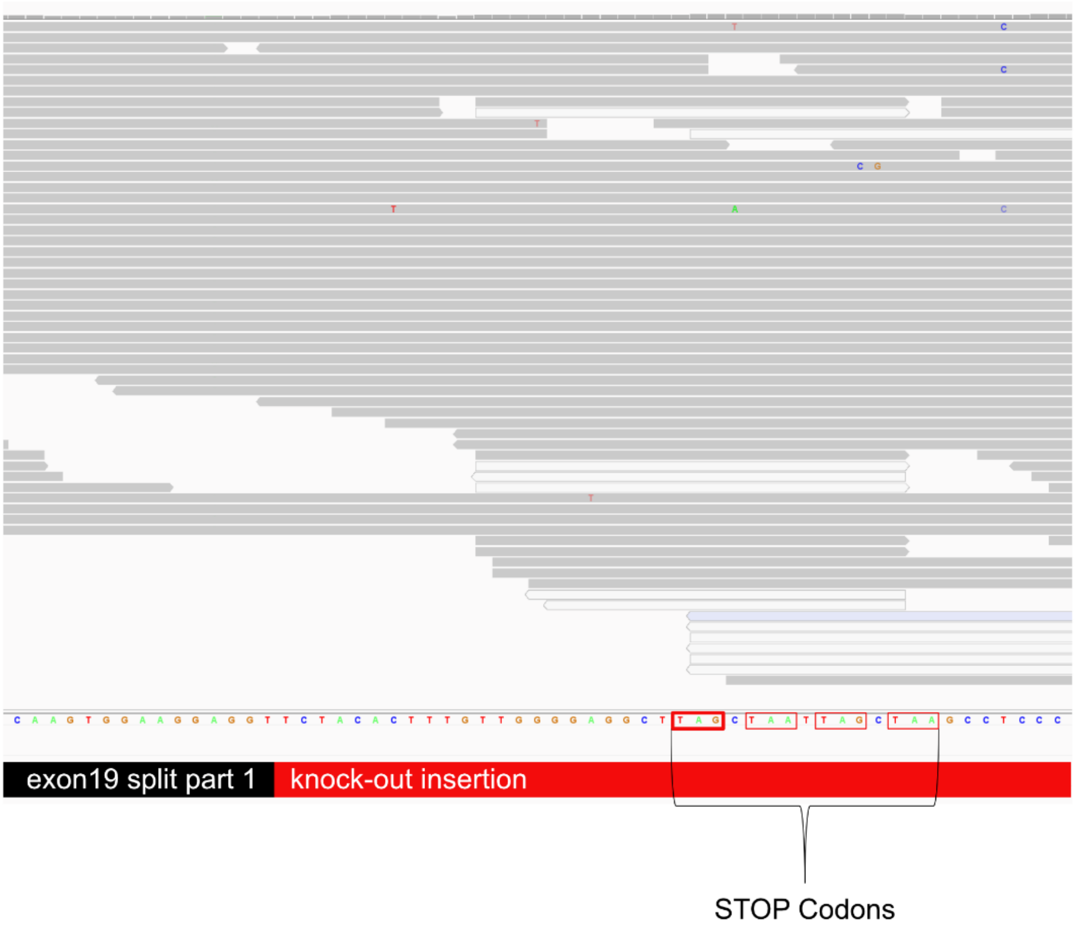


**Figure S4: Genomic characterization of the *Cr-kea1* CRISPR/Cas9 knock-out site (part2).** Integrative Genomics Viewer (IGV) visualization showing the left border at the insertion site within exon 19 of the *Cr-kea1* CRISPR line. Stop codons introduced via the mutation oligonucleotide are highlighted in red boxes.

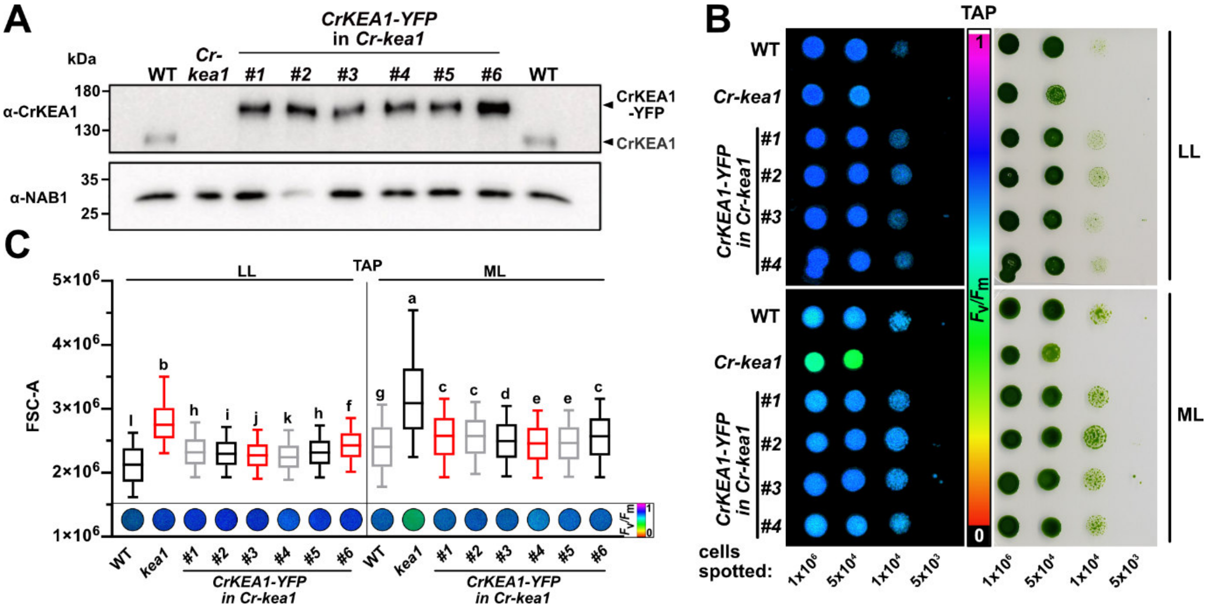


**Figure S5: Characterization of independent *CrKEA1–YFP* complementation lines. (A)** Immunoblot analysis of CrKEA1–YFP expression in independent complementation lines with *Cr-kea1* mutant background compared with wild type (WT) and the *Cr-kea1* mutant. Protein extracts were probed with anti-CrKEA1 antibodies. NAB1 served as a loading control. **(B)** Growth and photosynthetic performance of complementation lines on TAP drop cultures. Cultures were grown on solid TAP medium, treated for 4 days with the respective light intensity (LL, 25 µE; ML, 100 µE) and imaged using an Imaging-PAM system to visualize $F_v/F_m$. **(C)** Physiological characterization of complementation lines in liquid TAP cultures. Cultures pre-grown at 25 µE were exposed to either low light (LL, 25 µE) or moderate light (ML, 100 µE) for 24 h. Cell size distributions were determined by flow cytometry (FC), and maximum PSII quantum efficiency ($F_v/F_m$) was measured by PAM for the same samples. All experiments were performed under mixotrophic conditions in TAP medium. WT and *Cr-kea1* are included as controls. Statistical analysis was performed using a Kruskal–Wallis test with Dunn's post hoc test; n=3 replicates with > 10,000 cells were combined for the analysis.

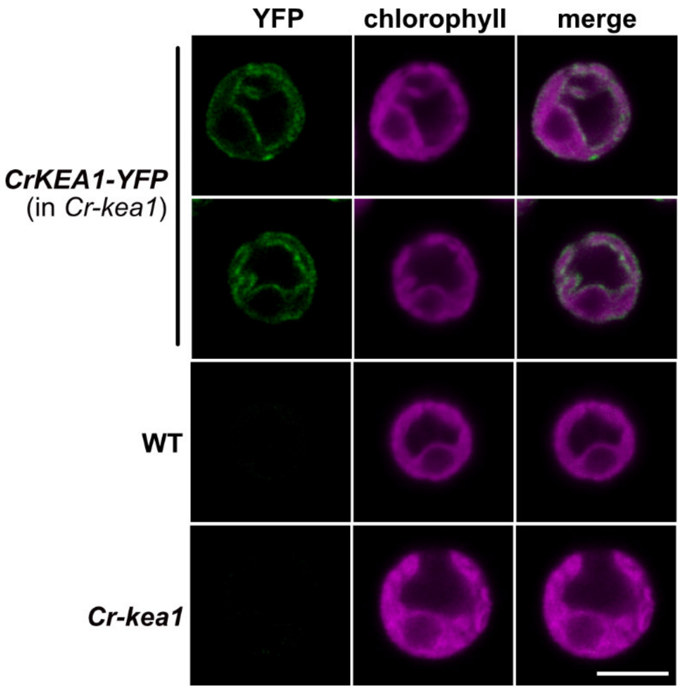


**Figure S6: CrKEA1-YFP localizes to the plastid envelope.** Confocal fluorescence microscopy showing plastid inner envelope localization of CrKEA1-YFP compared to control cells of the *Cr-kea1* background and wild type (WT) grown in TAP medium at 25 µE. Scale bar = 5 µm.

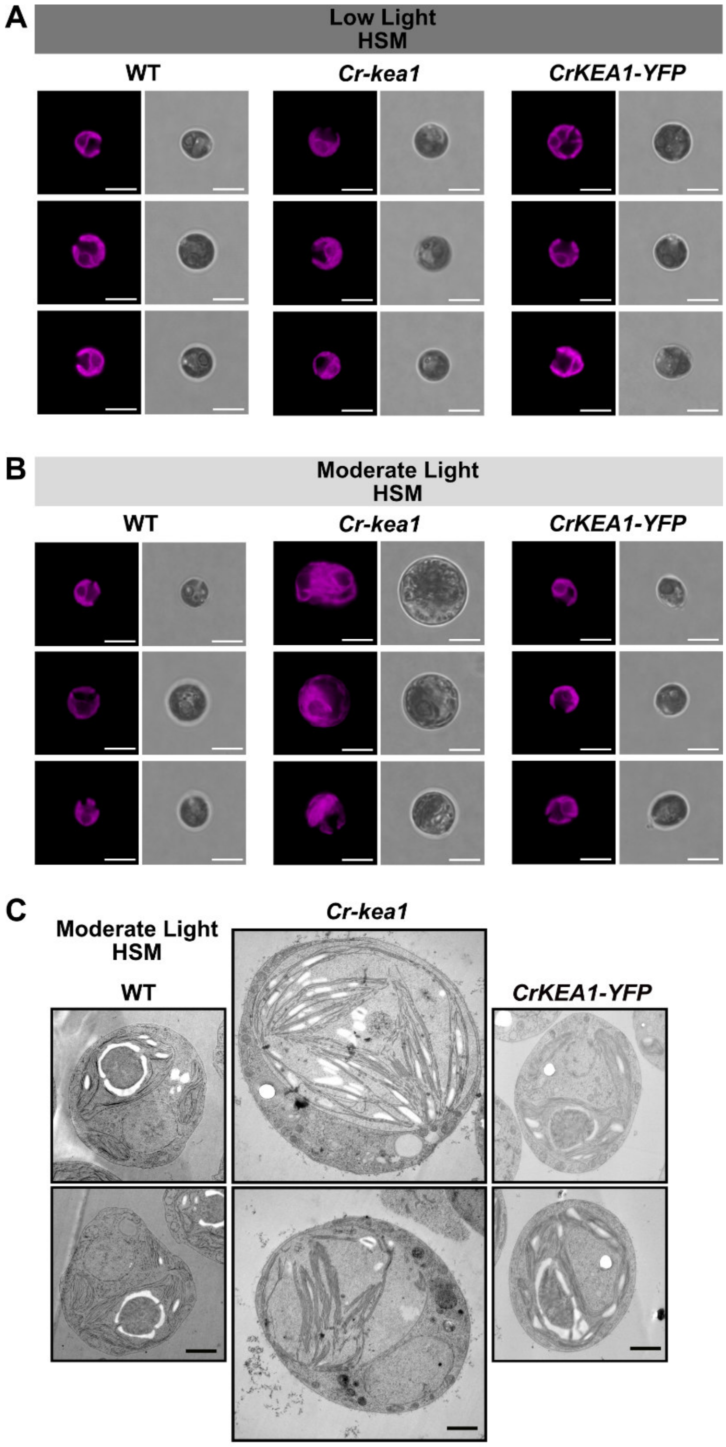


**Figure S7: Confocal microscopy of chloroplast morphology and ultrastructural analysis of *Cr-kea1* cells.** **(A)** Confocal microscopy of WT, *Cr-kea1*, and the *CrKEA1-YFP* complementation line grown in HSM medium under low light (LL, 25 µE) or moderate light (ML, 100 µE). Chlorophyll autofluorescence was used to visualize chloroplast morphology. Scale bar = 5 µm. **(B)** Additional transmission electron microscopy (TEM) images of WT and *Cr-kea1* cells grown in 100 µE in HSM corresponding to the ultrastructural analysis shown in Figure 1. Scale bar = 1 µm.

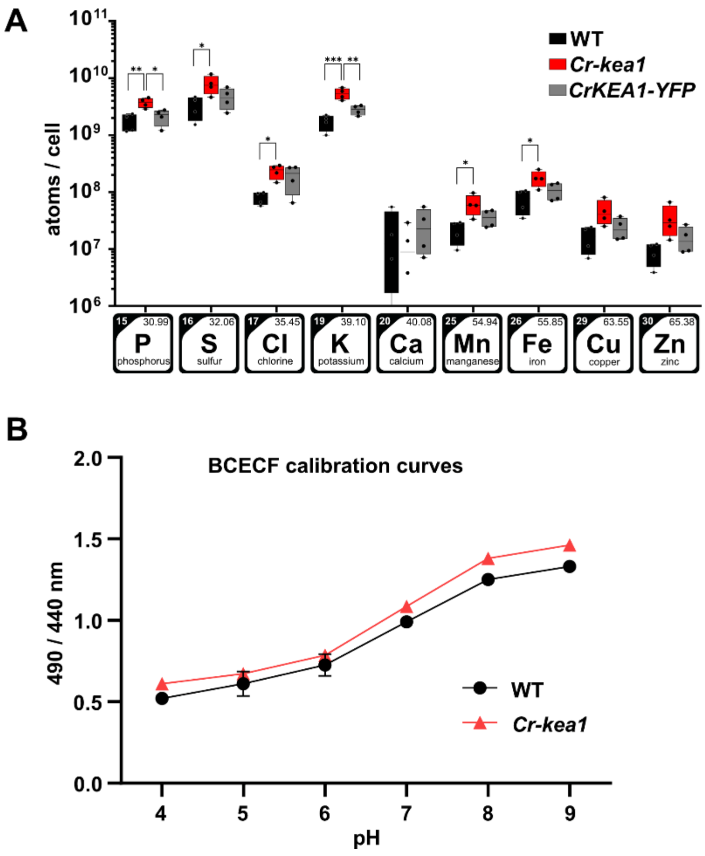


**Figure S8: Ionomic profiles expressed per cell and calibration of cytosolic pH measurements in *Cr-kea1*. (A)** Elemental composition of whole cells was quantified by Total Reflection X-Ray Fluorescence (TXRF) from TAP liquid cultures grown at 25 µE. Elemental abundances are expressed as atoms per cell to allow comparison with previously reported Chlamydomonas ionomic datasets. Four biological replicate cultures per genotype were analyzed. Data were evaluated by one-way ANOVA for each element separately, followed by Tukey's multiple comparison test; n=4. Box plots display the median (center line), the 25th–75th percentiles (box), and the minimum and maximum values (whiskers). Asterisks indicate significant differences (* $p < 0.05$; ** $p < 0.01$; *** $p < 0.001$). **(B)** Calibration curves for cytosolic pH determination using the ratiometric fluorescent dye BCECF. BCECF-loaded cells were equilibrated with buffers of defined pH (pH 4–9) in the presence of nigericin to equalize cytosolic and external pH. Fluorescence ratios (excitation 490/440 nm, emission 535nm) were recorded and plotted against buffer pH to generate standard curves used for cytosolic pH calculations. Calibration curves were determined separately for each genotype. n = 3.

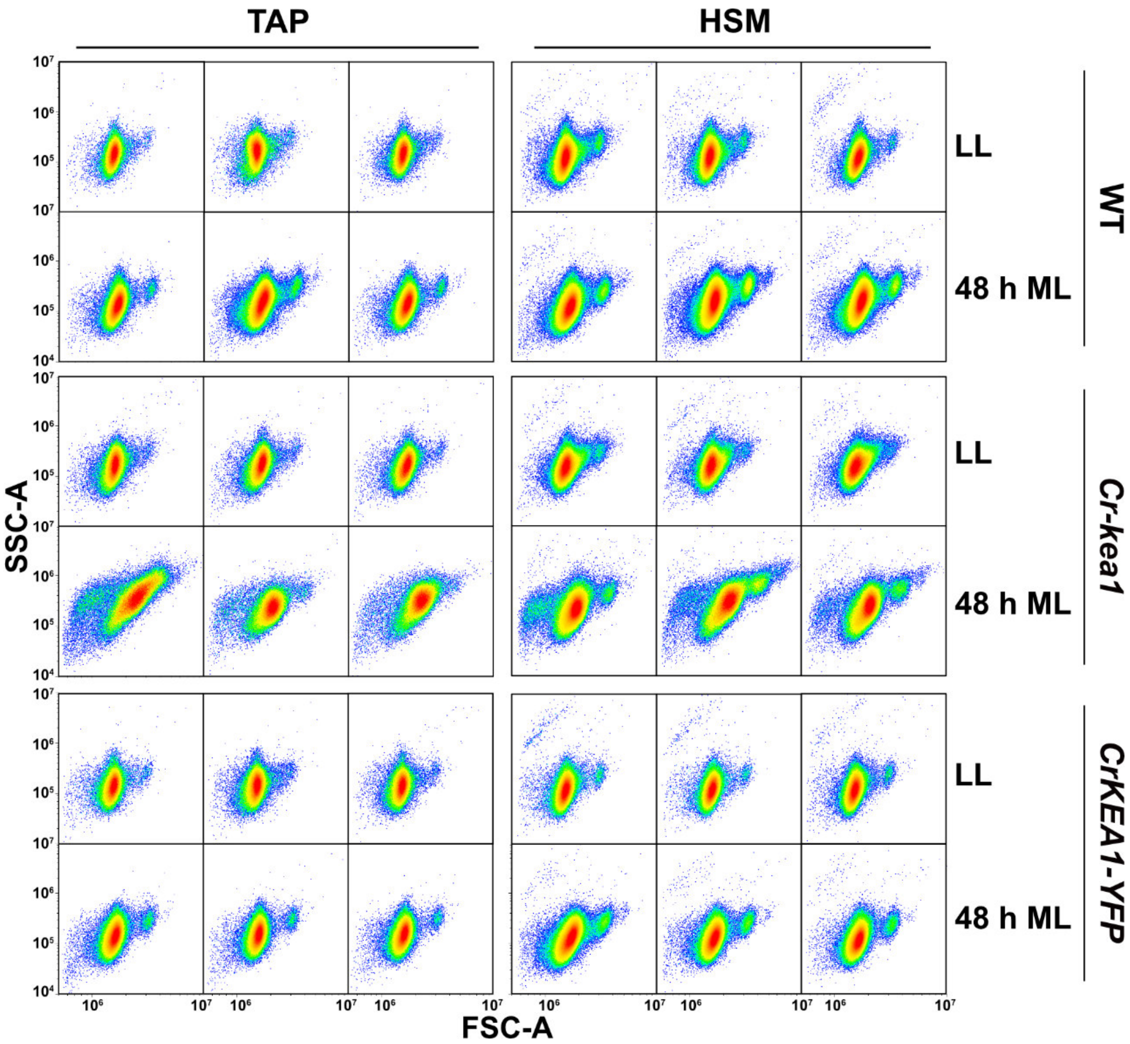


**Figure S9: Scatter plots corresponding to the flow cytometry experiment shown in Figure 3.**

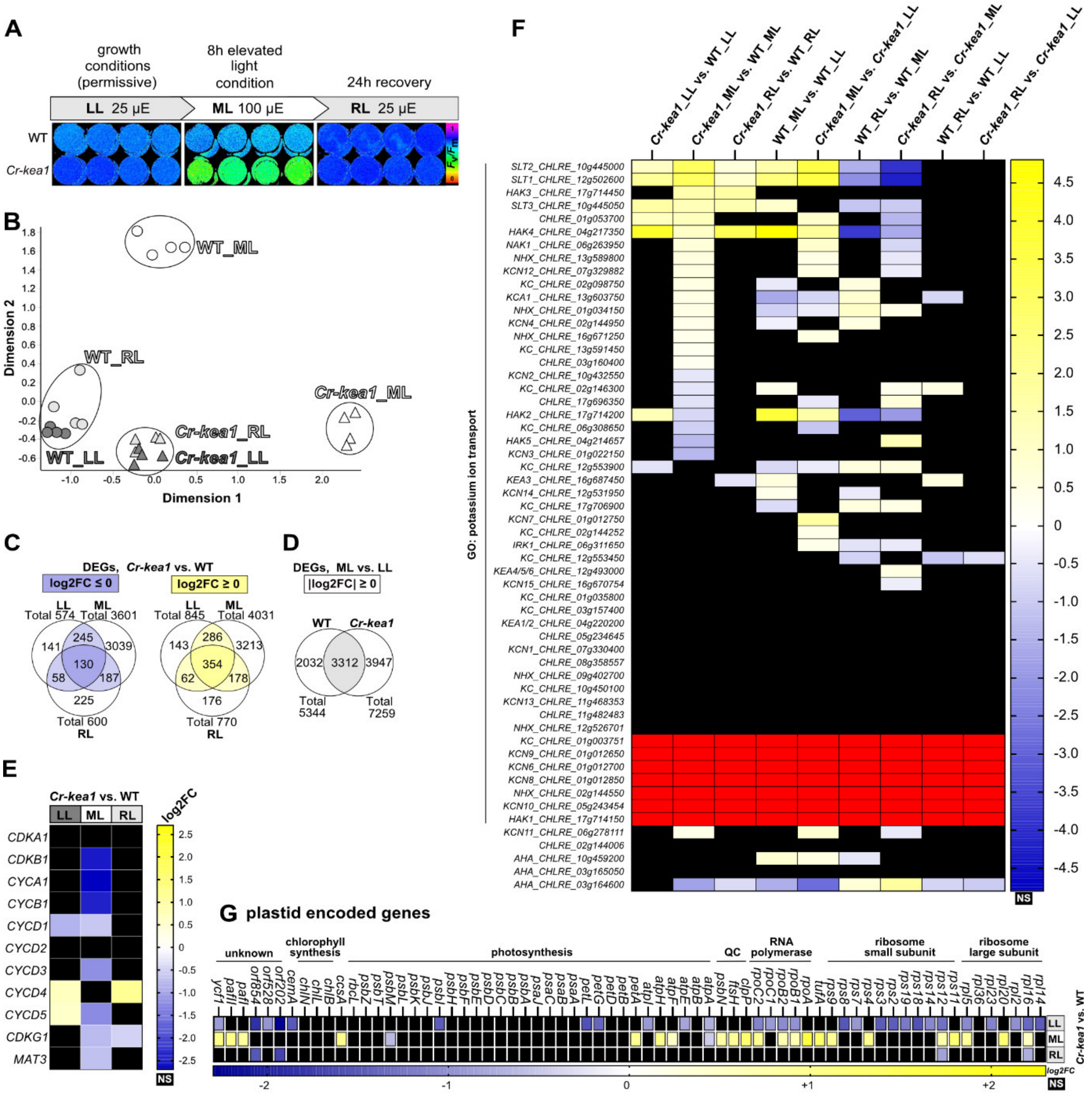

**Figure S10: Additional results related to the RNA-seq experiment. (A)** PAM fluorometry ($F_v/F_m$) images for all four biological replicates per condition and genotype. **(B)** Multidimensional scaling (MDS) plot showing the variance structure and clustering of RNA-seq replicates. Replicates of the same genotype and treatment cluster tightly, confirming data consistency. **(C)** Venn diagrams showing the overlap of significantly (adj. p-value < 0.05) upregulated or downregulated genes in *Cr-kea1* compared to WT under the different light conditions. **(D)** Venn diagram showing differentially expressed genes (DEGs) unique to *Cr-kea1* or shared WT after the light shift from low to moderate light, illustrating the genotype-specific transcriptional response to the light increase from LL to ML. **(E)** Heatmap showing $log_2FC$ values of selected DEGs related to the cell cycle. Not significant (NS) DEGs in black. **(F)** Heatmap showing $log_2FC$ values of selected DEGs encoding potassium transporters, ion channels, P-type ATPases, and other ion homeostasis related genes. Not significant (NS) DEGs in black. Genes without reads in red. **(G)** Heatmap showing $log_2FC$ values of plastid encoded genes. Not significant (NS) DEGs in black.

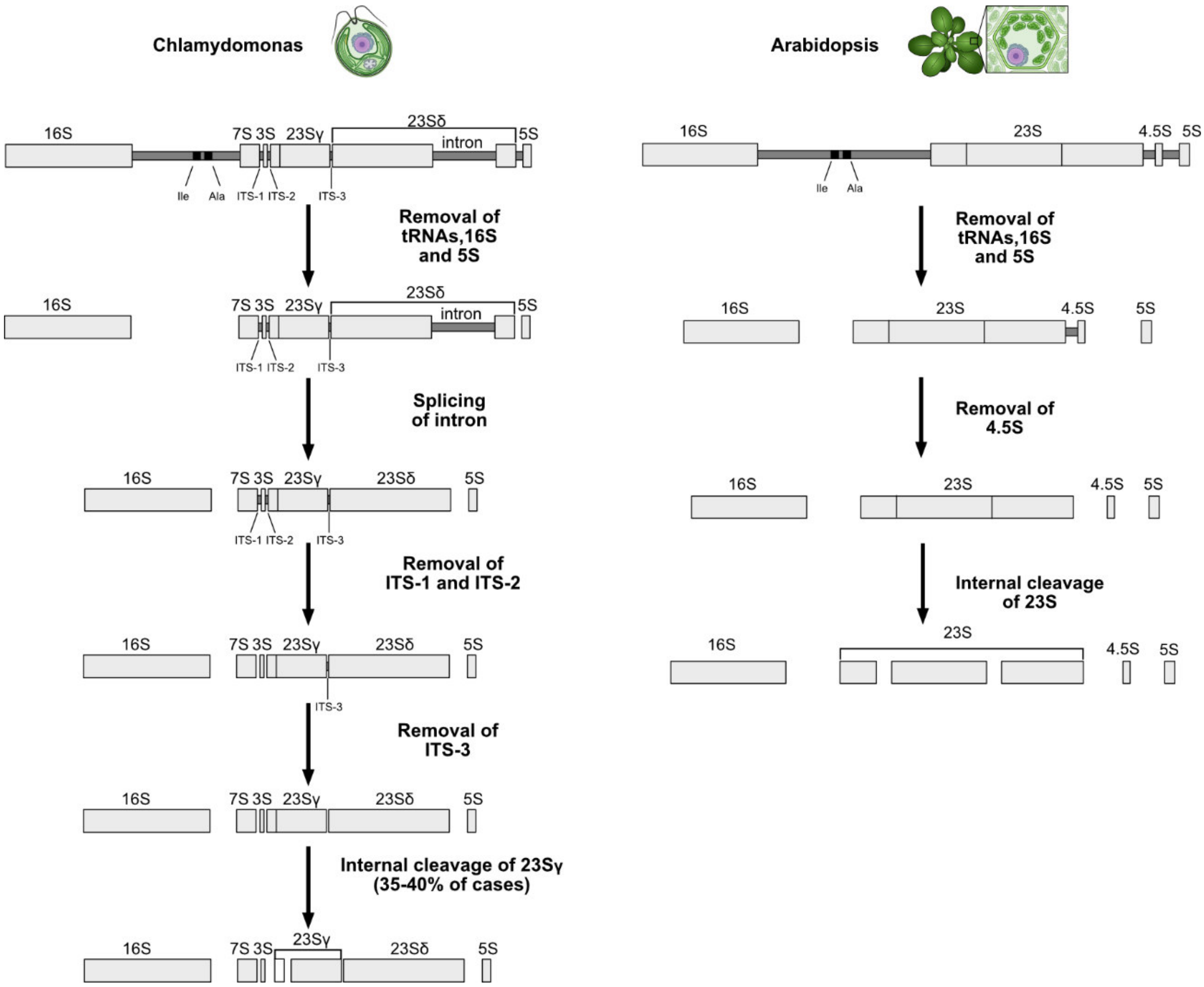


**Figure S11: Comparison of plastid rRNA processing pathways in Chlamydomonas and Arabidopsis.** Schematic representation of plastid rRNA cleavage and maturation pathways, illustrating the processing of precursor transcripts into the mature 16S and 23S rRNAs and their derived fragments. In both species, a 5S rRNA fragment is cleaved from the precursor; however, Arabidopsis uniquely generates a 4.5S fragment, whereas Chlamydomonas produces distinct 7S and 3S fragments from the 23S precursor. The schemata are adapted from Holloway and Herrin (1998) and Nishimura et al. (2010).

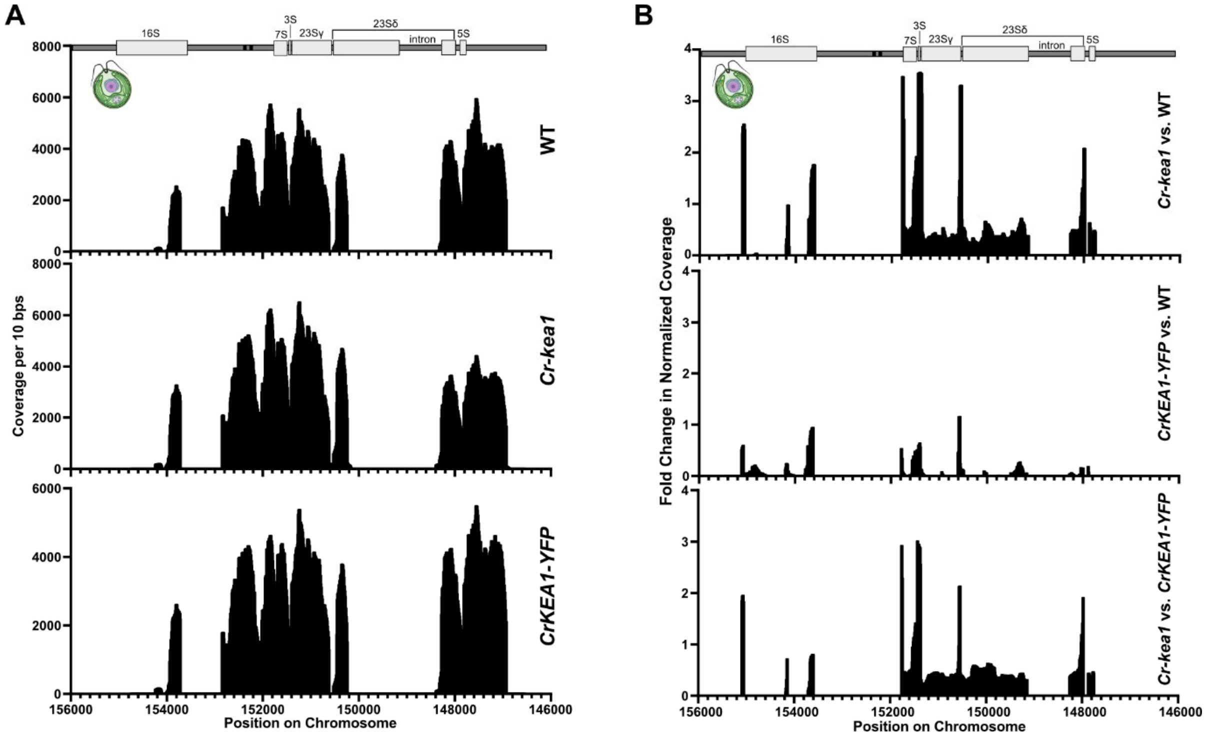


**Figure S12: rRNA-seq analysis of plastid rRNA processing including the *CrKEA1-YFP* complementation line. (A)** Read coverage profiles across the plastid rRNA operon for Chlamydomonas WT, *Cr-kea1*, and the *CrKEA1-YFP* complementation line. **(B)** Fold-change in normalized coverage (*BamCompare*) between *Cr-kea1* and WT, highlighting regions of precursor accumulation. The upper panel corresponds to the dataset shown in the main figures.

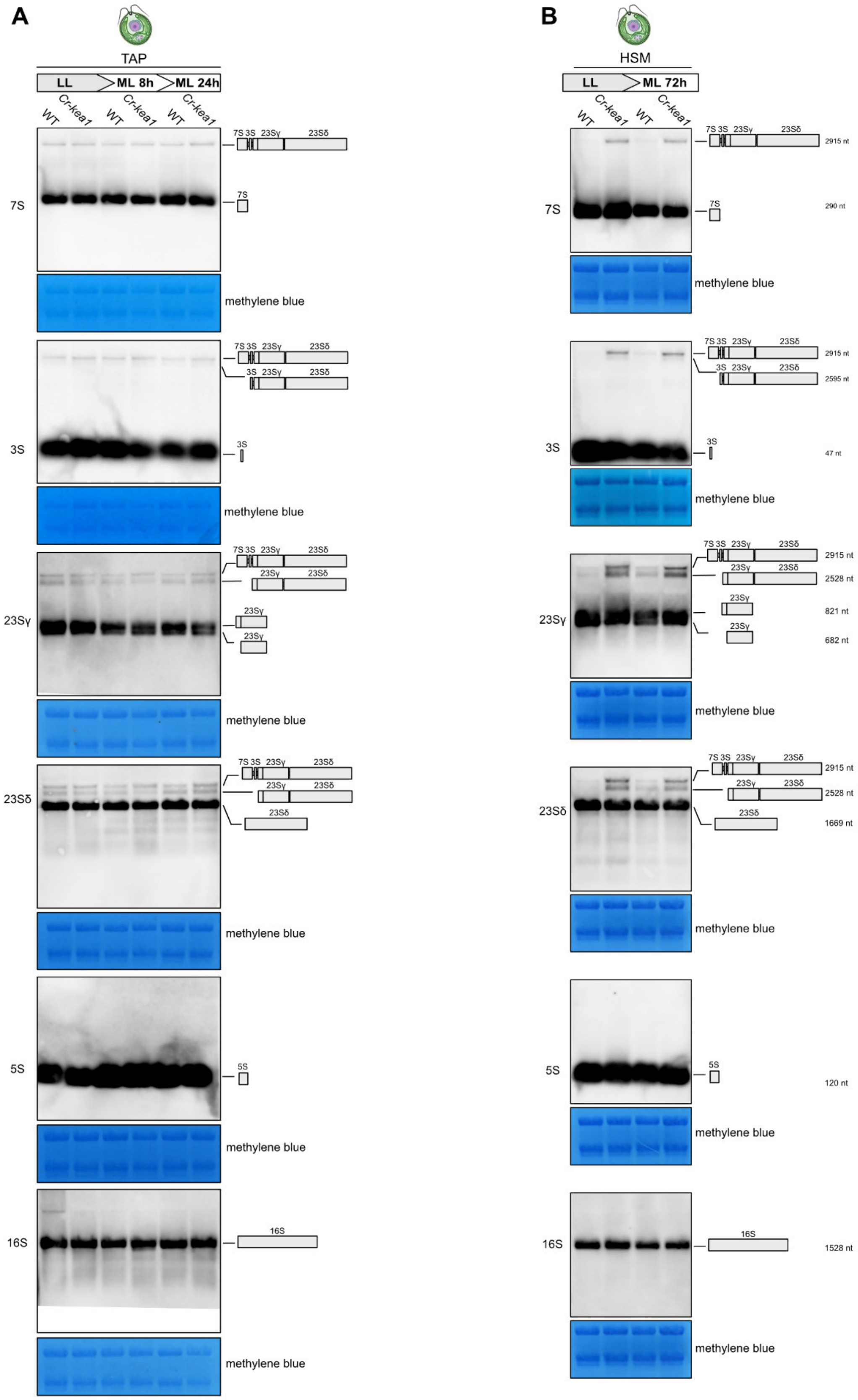


**Figure S13: RNA blot analysis of plastid rRNA processing in the *Cr-kea1* mutant.** RNA blots of Chlamydomonas cultures subjected to light-shift experiments using specific plastid rRNA probes show accumulation of the 7S–3S–23S precursor in *Cr-kea1* compared to WT. Cultures were grown in mixotrophic in TAP **(A)** or photoautotrophic in HSM **(B)** medium under low light (LL, 25 µE) and shifted to moderate light (ML, 100 µE). RNA loading was normalized to total RNA.

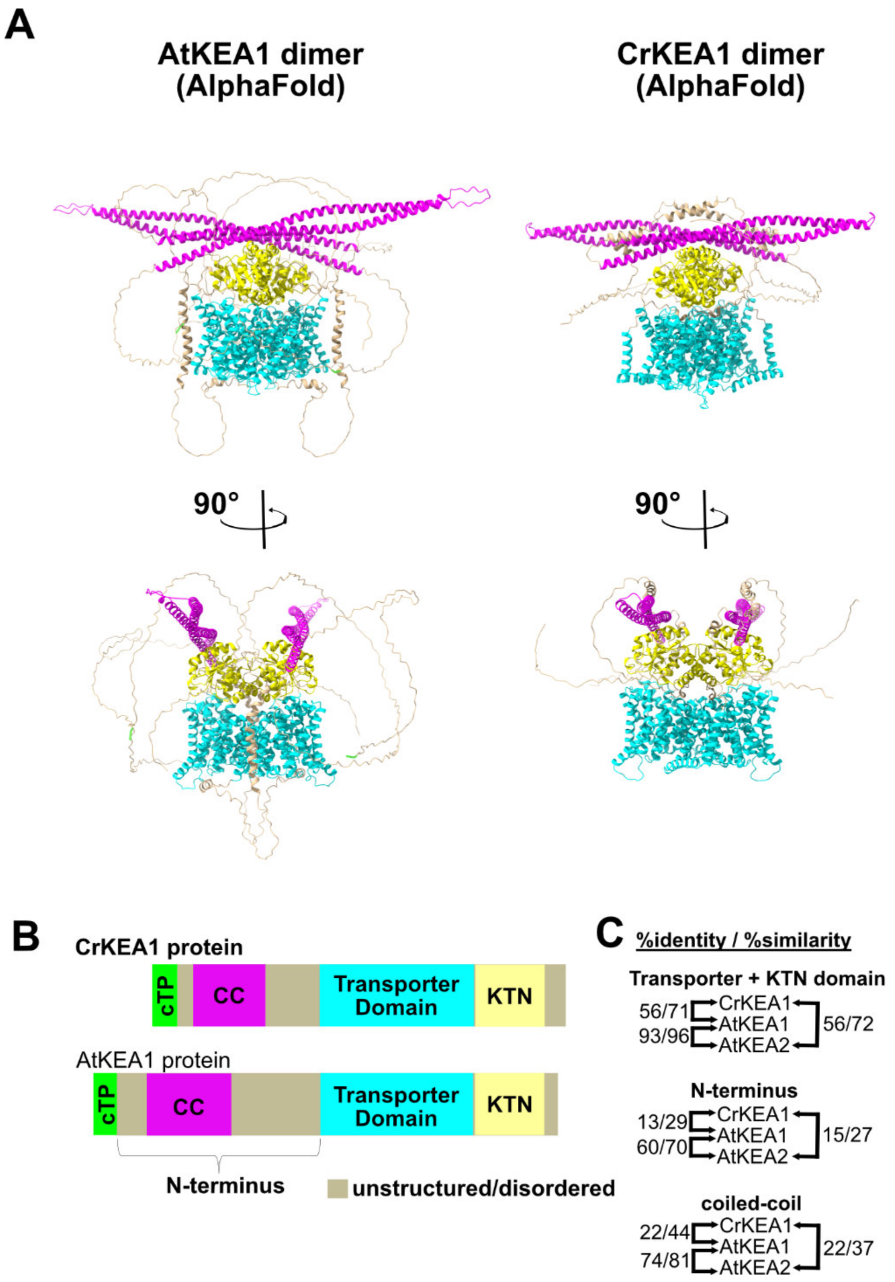


**Figure S14: Structural comparison of AtKEA1 and CrKEA1. (A)** AlphaFold3-predicted dimeric models of Arabidopsis KEA1 (AtKEA1) and its Chlamydomonas homolog (CrKEA1). Color coding corresponds to the domain organization shown in panel (B). Dimer selection was guided by the structural findings for the bacterial homolog KefC (Gulati et al., 2024). **(B)** Simplified domain architecture of AtKEA1 and CrKEA1. **(C)** Sequence identity and similarity scores between AtKEA1, AtKEA2, and CrKEA1.

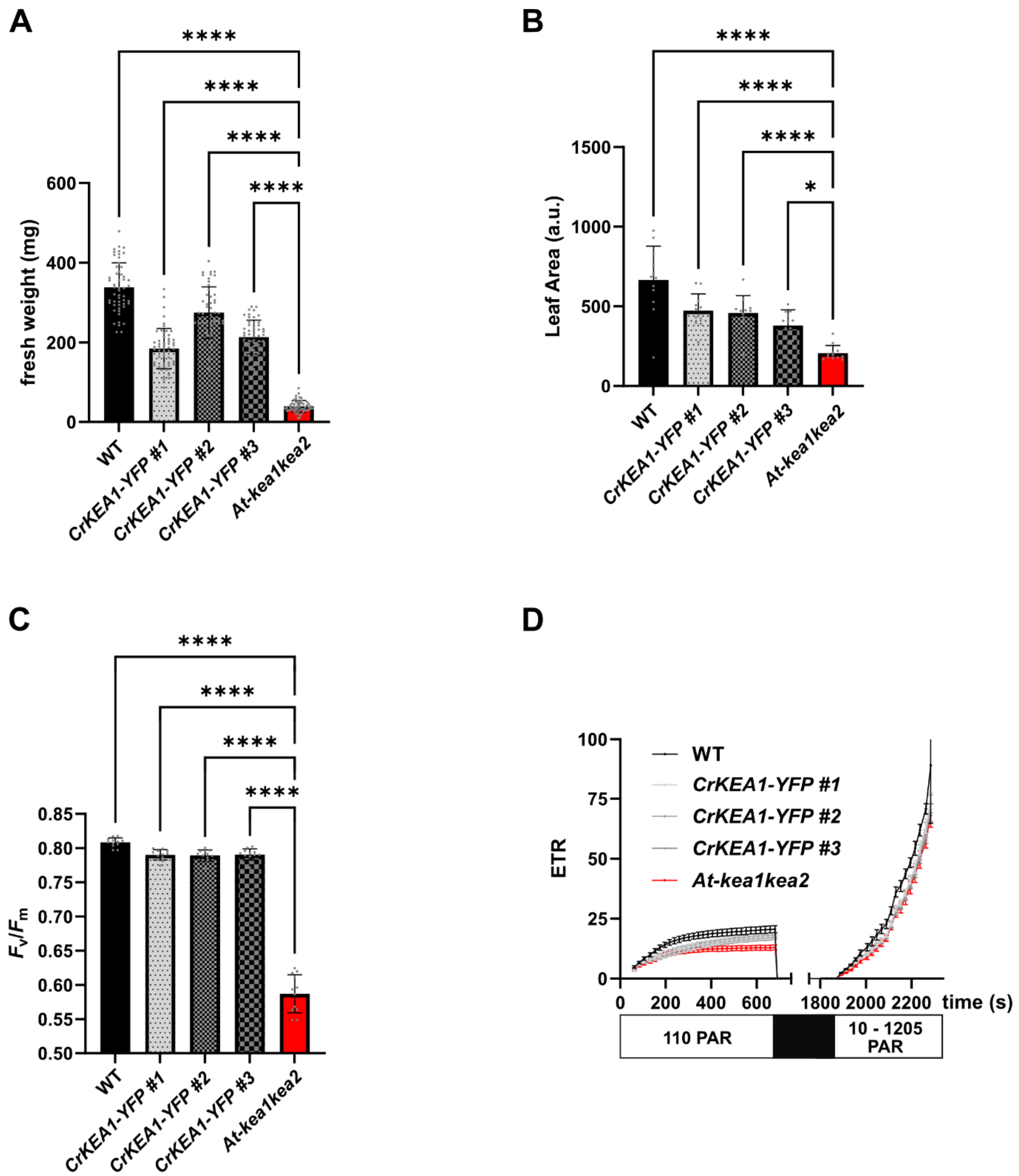


**Figure S15: Characterization of the Arabidopsis *At-kea1kea2* line complemented with CrKEA1-YFP. (A)** Rosette fresh weight (n=50) **(B)** Leaf area (n=12). **(C)** Maximum quantum yield of PSII ($F_v/F_m$) (n=12). **(D)** Electron transport rate (ETR) (n=12). Measurements were performed under a light curve protocol with 11 min at 110 PAR, followed by 20 min dark adaptation. Subsequently, light intensity was increased in 20 s intervals through the following PAR levels: 10, 20, 35, 55, 80, 110, 145, 185, 230, 395, 485, 530, 610, 700, 800, 925, 1075, and 1250 PAR. Mean±SD and one-way ANOVA followed by Dunnett's multiple comparisons test is shown. Statistical significance is indicated by asterisks (* $p < 0.05$; ** $p < 0.01$; *** $p < 0.001$; **** $p < 0.0001$).

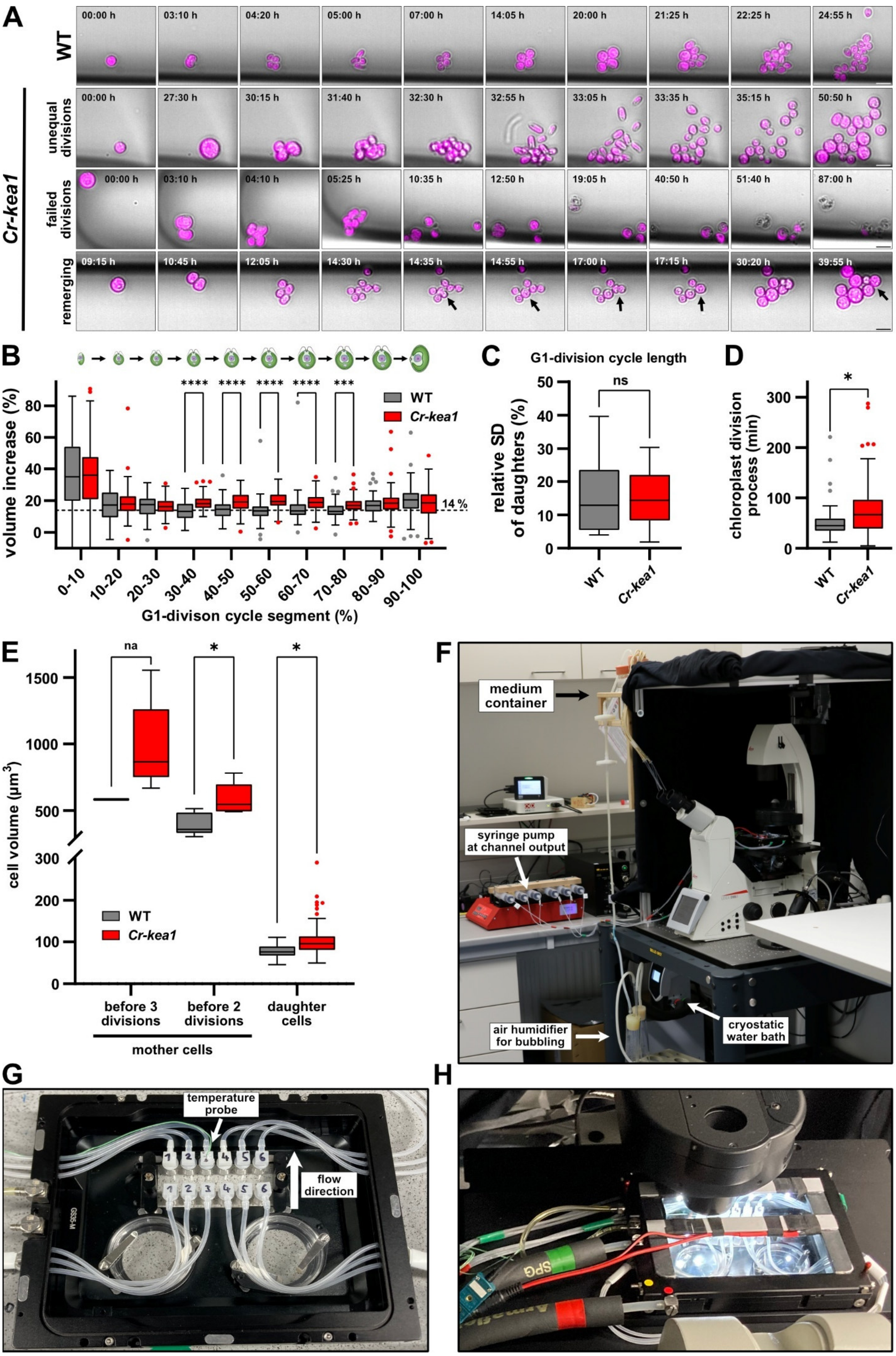
A
WT
Cr-kea1
unequal divisions
failed divisions
remerging
00:00 h
03:10 h
04:20 h
05:00 h
07:00 h
14:05 h
20:00 h
21:25 h
22:25 h
24:55 h
27:30 h
30:15 h
31:40 h
32:30 h
32:55 h
33:05 h
33:35 h
35:15 h
50:50 h
04:10 h
05:25 h
10:35 h
12:50 h
19:05 h
40:50 h
51:40 h
87:00 h
09:15 h
10:45 h
12:05 h
14:30 h
14:35 h
14:55 h
17:00 h
17:15 h
30:20 h
39:55 h
B
volume increase (%)
G1-divison cycle segment (%)
0-10
10-20
20-30
30-40
40-50
50-60
60-70
70-80
80-90
90-100
14 %
WT
Cr-kea1
C
G1-division cycle length
relative SD of daughters (%)
ns
D
chloroplast division process (min)
E
cell volume (µm³)
na
before 3 divisions
before 2 divisions
daughter cells
mother cells
F
medium container
syringe pump at channel output
air humidifier for bubbling
cryostatic water bath
G
temperature probe
flow direction
H

**Figure S16: Single cell live imaging of *Cr-kea1* and WT. (A)** Extended version of time lapse imaging shown in the main figure. Scale bar, 10 µm. **(B)** Quantitative analysis of single-cell trajectories with focus on the volume increase during the G1-division cycle. **(C)** Variation of the G1-division cycle length of WT and *Cr-kea1* daughter cells. $n^{WT}=68$; $n^{kea1}=90$. **(D)** Chloroplast division time in WT and *Cr-kea1* mutants. $n^{WT}=16$; $n^{kea1}=16$. **(E)** Volume of mother cells before undergoing 3 ($n^{WT}=1$; $n^{kea1}=12$) or 2 divisions ($n^{WT}=14$; $n^{kea1}=5$) and volume of the resulting daughters ($n^{WT}=63$; $n^{kea1}=88$). **(F)** Imaging setup for long-term single-cell recordings on a Leica DMi8 Thunder microscope. Bubbled medium was gravity fed into hydrogel channels. Flow was controlled by a syringe pump (KF Technology, NE-1600) at the channel output set to 200 µL $h^{-1}$ in suction mode. Temperature was maintained at 23°C using a cryostatic water bath (Lauda, RE415S). An air humidifier was used for medium bubbling to prevent evaporation and resulting osmolarity changes. **(G)** Hydrogel-based microfluidic flow-cell chip (based on Ibidi µ-Slide VI 0.4) equipped with a temperature probe and connected tubing for continuous medium perfusion. **(H)** Flow-cell channels positioned inside a stage-top heating and cooling chamber (Okolab, H101-CRYO-BL) with LED illumination for controlled light exposure during imaging. *For all panels:* Box plots follow Tukey's convention. Statistical significance was determined with Mann-Whitney tests and is indicated by asterisks (* $p < 0.05$; ** $p < 0.01$; *** $p < 0.001$; **** $p < 0.0001$).
